# Coordinated control of both microtubule ends regulates mitotic spindle length

**DOI:** 10.64898/2026.02.08.704347

**Authors:** Shane A. Fiorenza, Sheba Cheeran, Elena Doria, Iva M. Tolić, Patrick Meraldi, Nenad Pavin

## Abstract

The mitotic spindle is a biomechanical structure whose length must be precisely controlled to ensure faithful chromosome segregation. The treadmilling of microtubules towards centrosomes, termed poleward flux, is involved in spindle length control. However, poleward flux has been shown to be both inversely and directly proportional to spindle length, a contradiction that remains unexplained by current mechanisms. Here we introduce a model which demonstrates that length-dependent regulation at both microtubule ends allows plus- and minus-end dynamics to synchronize with one another, enabling pathways of poleward flux-based spindle length control which can rectify previous results. Moreover, our model predicts that spindle length and poleward flux can vary independently via simultaneous perturbations at both microtubule ends, which we experimentally validate with combinations of KIF18A, KIF2A, and KATNB1 depletions in human cells. Our results thus resolve a longstanding paradox and provide mechanistic insight into the reciprocal control of spindle length and poleward flux.

---

A fundamental prerequisite for the successful propagation of life is the ability for cells to divide and pass on their genetic material to the next generation. To facilitate this process and minimize any potential errors, cells employ the mitotic spindle, a micromachine composed of microtubules and associated proteins.^1^ The spindle generates forces on chromosomes that are regulated over both space and time to ensure a high level of fidelity throughout mitosis.^2,3^ Due to its biological importance, the mechanisms controlling spindle length and function have been extensively studied both *in vivo* and *in vitro*.^4–18^ Given the complexity of the mitotic spindle and experimental resolution limits, theoretical modeling has also been a powerful tool in understanding how the spindle sets its length^4–6,19–23^ and shape.^24,25^ However, the mitotic spindle utilizes and precisely coordinates a myriad of feedback loops and regulatory processes which are difficult to elucidate the individual contributions of, and so our understanding of the physical mechanisms responsible for spindle length regulation remains incomplete.

In metazoan spindles, the continual depolymerization of microtubule minus-ends at centrosomes leads to a net velocity towards the spindle pole referred to as poleward flux, ^26–30^ and microtubules maintain a constant length via compensatory polymerization at plus-ends. Poleward flux velocity is precisely modulated throughout mitosis^31–33^ and is important for facilitating chromosome movement^34–36^ and positioning^37^ as well as preventing erroneous kinetochore attachments.^38^ Spindle length is also thought to be regulated by poleward flux,^39–41^ but its role is obfuscated by seemingly contradictory experiments in which reducing poleward flux leads to spindle length decreasing,^40,42–45^ increasing,^34,41,46^ or remaining approximately unchanged.^35,47^ Additionally, poleward flux velocity can span over an order of magnitude between species, e.g., 0.5 um/min in LLC-PK1 cells,^29^ 1.3 um/min in humans,^43^ and 5.2 um/min in Drosophila embryos,^31^ despite comparable spindle lengths of 18.8, 13.9, and 10-14 microns, respectively. This suggests that cells can tolerate vastly different poleward flux velocities and maintain their spindle length relatively independently of the quantity.

Theoretical models have been used to study how poleward flux is involved in chromosome movement, positioning, and error-correction.^43,48–52^ In these models, depolymerization at microtubule minus-ends and pulling at the centrosome leads to poleward movement which generates forces on kinetochores. Forces at the centrosomes are balanced by either polar ejection forces^48,49,51^ or motor sliding forces^43,52^ or are not considered in the model.^50^ The direct relationship between spindle length and poleward flux velocity has also been studied theoretically, expanding upon these previous works by incorporating a length-dependent depolymerization rate at microtubule minus-ends.^23^ However, experimental data in which spindle length and poleward flux exhibit an inverse relationship or are independent of each other are still unexplained. Therefore, how poleward flux regulates spindle length despite substantially varying values and inconsistent proportionality remains an open question.

In this paper, we use a combination of theory and experiment to explore how mitotic spindle length is regulated by poleward flux. We introduce an analytic model that incorporates length-dependent mechanisms at both the plus- and minus-end of microtubules in addition to core ingredients such as motor sliding and interkinetochore forces. We find that this combination of length-dependent mechanisms allows each microtubule end to sense and respond to changes at the opposite end, resulting in mutual synchronization. Consequently, perturbations of plus- and minus-end dynamics influence poleward flux velocity in opposite ways, allowing our model to explain both direct and inverse relationships between poleward flux and spindle length. Our model also predicts that by perturbing both ends of the microtubule simultaneously, these changes can cancel each other out, allowing poleward flux and mitotic spindle length to vary independently of one another. We confirm these predictions with *in vivo* depletion experiments, which show that co-depletion of the plus-end polymerization inhibitor KIF18A and the minus-end severase KATNB1 can rescue poleward flux velocity for larger spindles. In summary, our results demonstrate that length-dependent mechanisms at both microtubule ends allow plus- and minus-end dynamics to synchronize with one another by communicating through microtubule length, facilitating robust spindle length control over a wide range of poleward flux velocities.

## A model incorporating length-dependent mechanisms at both microtubule ends predicts mutual synchronization of plus- and minus-end dynamics

To investigate mechanisms of mitotic spindle length regulation, we introduce a model which includes length-dependent mechanisms at both the plus- and minus-end of microtubules (Fig. 1, Methods). The central idea of our model is to study the potential interactions between these two length-dependent mechanisms and explore how they influence the relationship between poleward flux and spindle length. The model includes a pair of microtubule fibers originating from opposite centrosomes that are bound to sister kinetochores, referred to as *k-fibers*, and a pair of microtubule fibers that interdigitate with other microtubules to form anti-parallel overlaps, referred to as *bridging fibers*.^53^ We use the antenna model to describe the length-dependent regulation of microtubule plus-ends^54^ (Fig. 1, inset 3) and extend it to microtubule minus-ends as well (Fig. 1, inset 1). Length-dependent plus-end regulation by proteins such as KIF18A is already well-established in the mitotic spindle,^54–57^ and we justify the extension to minus-ends with previous experiments which have shown that the abundance of minus-end proteins KATNB1 and KIF2A at centrosomes correlates with the length of microtubules emanating from it.^58^ Forces in the system are generated at k-fiber plus-ends by kinetochores and laterally along microtubules by plus-end-directed crosslinking motors that accumulate in anti-parallel overlaps (Fig. 1, inset 2), and these forces are transmitted to the centrosome through the minus-ends of microtubules. We keep overlap length fixed in our model because the mechanisms which regulate it are outside the scope of this paper and not well understood. This model provides a framework for studying the role of microtubule plus- and minus-end length regulators in poleward flux velocity and spindle length control.

**Figure 1.**
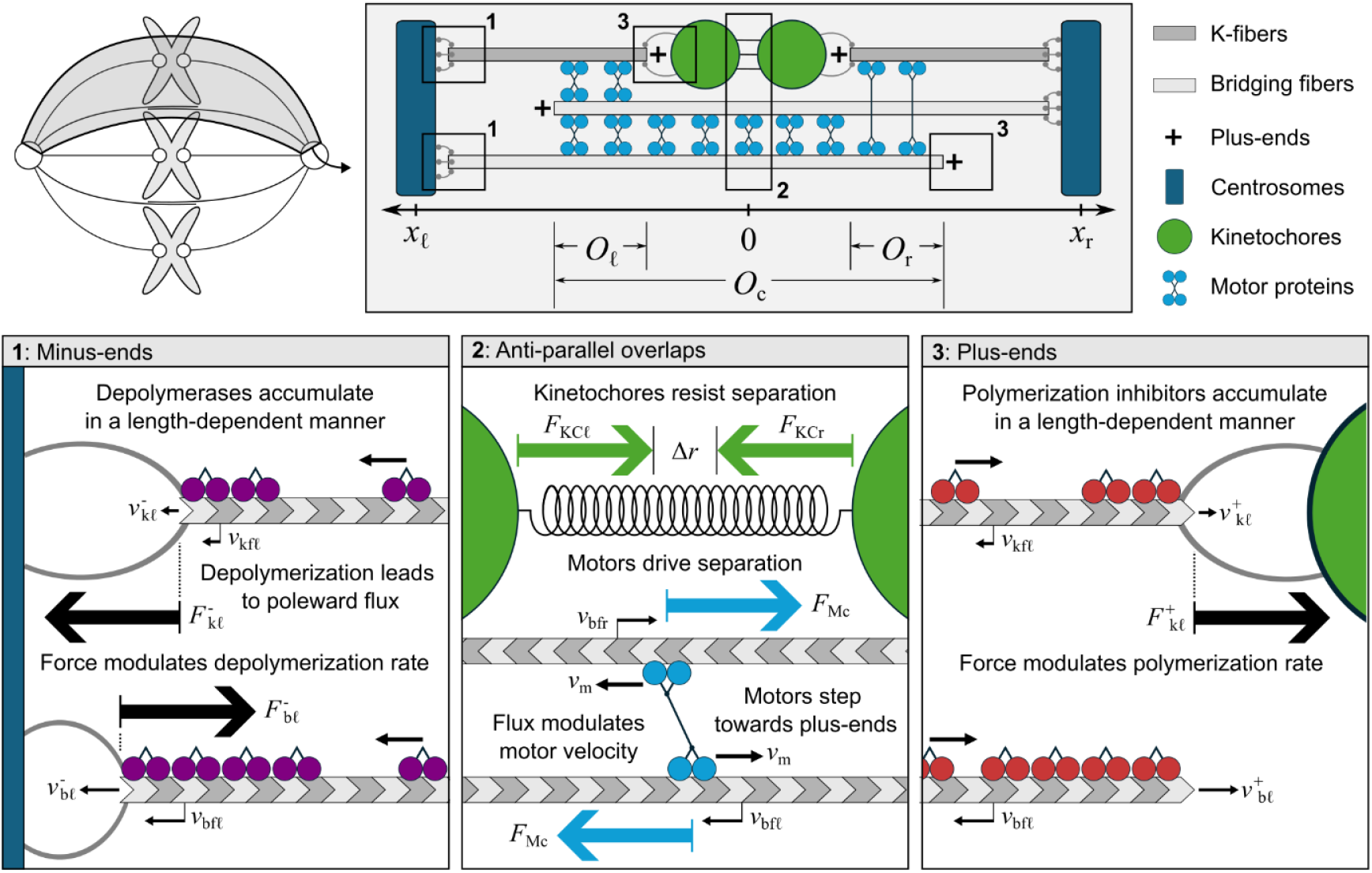
Model schematic. Our one-dimensional model represents a single chromosome pair out of the full mitotic spindle. Left and right centrosomes (blue) each have two microtubules (grey) attached to them. One microtubule is permanently affixed to the nearest sister kinetochore (green) to model k-fibers, while the other extends past the spindle midzone to model bridging fibers. This configuration leads to three distinct anti-parallel overlap regions: one central overlap, O_c_, and two smaller overlaps to the left and right, O_*ℓ*_ and O_r_, respectively. Motors (light blue) bind to these overlap regions and produce forces that work to extend the spindle by sliding microtubule plus-ends towards one another. The linkage between sister kinetochores provides a resistive force that opposes the sliding by motors. (inset 1) Scheme demonstrating minus-end behavior in our model. There is a constant depolymerization rate that is modulated by both force (large black arrows) and length-dependent accumulation of depolymerase proteins (purple). Depolymerization of microtubule minus-ends leads to poleward flux. (inset 2) Scheme demonstrating force generation. As motors step towards the plus-ends of microtubules in anti-parallel overlaps, they generate force (large blue arrows) that moves minus-ends apart. The interkinetochore force (large green arrows) resists motor forces. Motor stepping velocity depends on the poleward flux velocities of the two microtubules it is bound to. (inset 3) Scheme demonstrating plus-end behavior in our model. There is a constant polymerization rate that is modulated by both force (large black arrow) and length-dependent accumulation of regulating proteins (red). Bridging fibers are considered free, so no force is exerted on their plus-ends. Polymerization at plus-ends balances depolymerization at minus-ends to facilitate poleward flux.

We first compare the model with and without length-dependent mechanisms in order to elucidate the contribution of each component. In the version of our model with no length-dependent regulatory mechanisms, there are three key parameters that set spindle length: (i) baseline polymerization velocity at the plus-end, 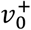, (ii) baseline depolymerization velocity at the minus-end, 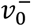 and (iii) motor sliding velocity, *υ*_0_. We begin with the simplest parameter choice: all three velocities are set equal, i.e., 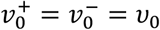. Under these conditions, the microtubule minus- and plus-ends remain at fixed positions, resulting in a constant spindle length (Fig. 2a). Next, we solve the case with faster baseline polymerization at plus-ends, i.e., 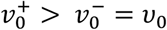, which results in the spindle length continually growing. Finally, we solve the model with slower baseline polymerization at plus-ends, i.e., 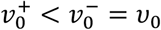. In this case, the centrosomes continually approach each other until the spindle collapses. Taken together, these results show that a model with no length-dependent mechanisms leads to a failed spindle which does not have a well-defined steady-state length and cannot withstand any perturbations (Fig. 2b). Therefore, our model suggests that some length-dependent mechanism must be present.

**Figure 2.**
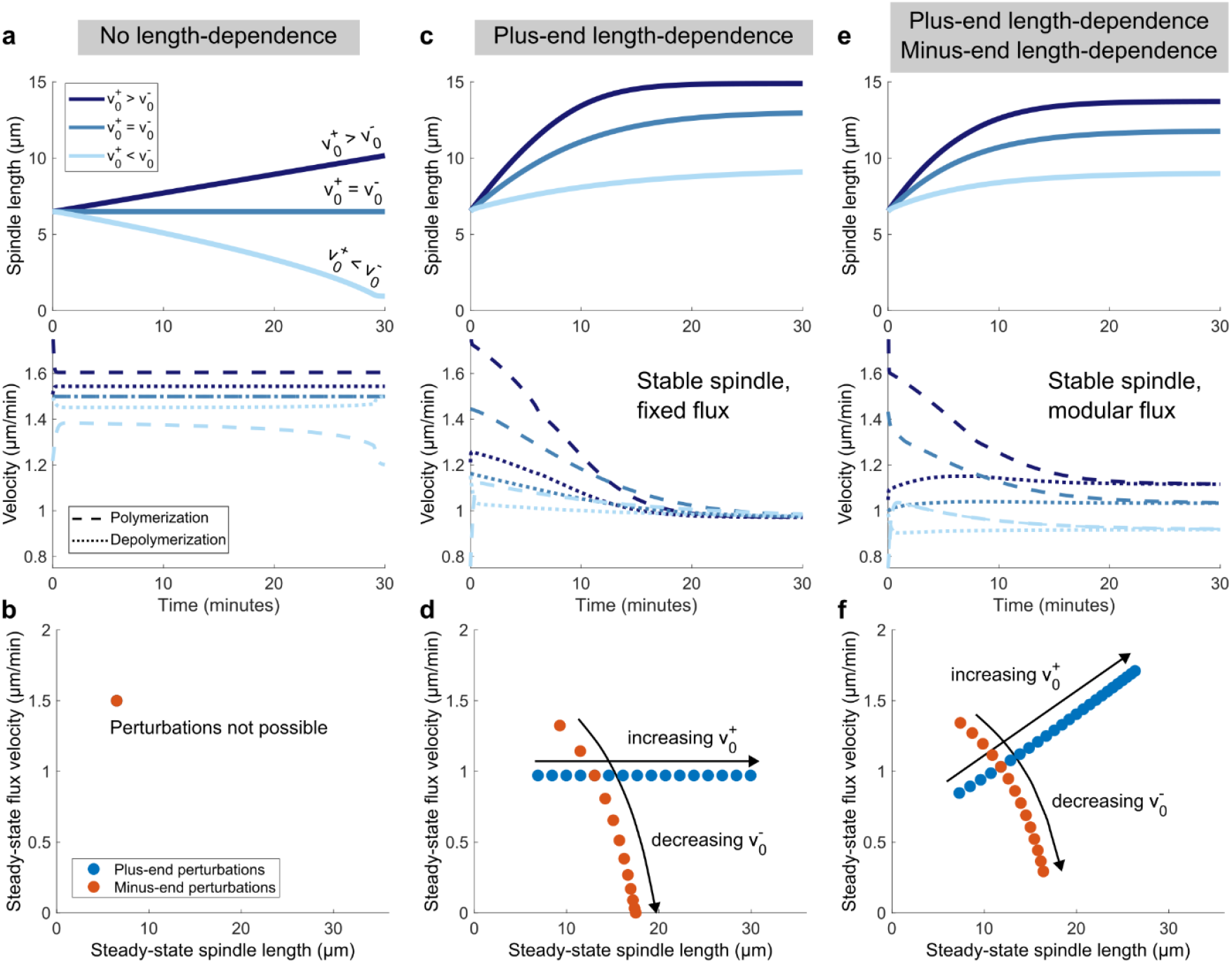
Length-dependent mechanisms at both microtubule ends facilitate the synchronization of plus- and minus-end dynamics. **a**, Plots of spindle length (top) and velocities of polymerization and depolymerization (bottom) over time for a version of the model that has no length-dependent mechanisms. Polymerization rate, depolymerization rate, and motor stepping velocity are all set to 1.5 μm/min to produce a constant geometry (blue). Increasing polymerization rate by 25% results in unbounded growth (dark blue, top) because polymerization is always larger than depolymerization (dark blue, bottom). Decreasing polymerization by 20% results in spindle collapse (light blue, top) because depolymerization is always larger than polymerization (light blue, bottom). **b**, Plot of steady-state spindle length and steady-state poleward flux velocity for different polymerization and depolymerization values, which only has one point since perturbations are not possible when no length-dependent mechanisms are present. **c** and **d**, Same plots as panels (**a**) and (**b**), but for a version of the model with a plus-end length-dependent mechanism present. In panel (**c**), perturbations correspond to polymerization rate being increased by 100% (dark blue) and decreased by 50% (light blue) relative to baseline (blue). All perturbations result in a stable spindle length (top), because the polymerization rates converge to the same depolymerization rate (bottom) in all cases. In panel (**d**), the plus-end scaling factor *A*^+^ and minus-end depolymerization rate *v*^−^ are varied from 10% to 500% of their nominal values, showing that plus-end perturbations are unable to influence flux when only a plus-end length-dependent mechanism is present. **e** and **f**, same plots as panels (**a**) and (**b**), but for a version of the model with both plus- and minus-end length-dependent mechanisms present. In panel (**e**), perturbations are the same as in (**c**). Spindle length is still stable for all perturbations (top), but now each polymerization rate converges to a different depolymerization rate (bottom). In panel (**d**), the plus-end scaling factor 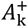 and minus-end scaling factor *A*^−^ are varied from 10% to 500% of their nominal values, showing that a length-dependent mechanism at the minus-end allows plus-end perturbations to influence poleward flux velocity. Parameters used for plots (**c**-**f**) appear in Table 1.

Plus-end length-dependent mechanisms are well established in the mitotic spindle,^54–57^ so we next explore a version of our model where only plus-end length-dependent mechanisms are included (Fig. 2c). We encode this mechanism into our model as a linear decrease in plus-end polymerization rate as microtubule length increases, becoming negative above a critical length to represent depolymerization outweighing polymerization for very long microtubules (Methods). With this mechanism added, we see that the spindle now reaches a well-defined steady state for each value of polymerization velocity used (Fig. 2c, top). This occurs because polymerization rates are modulated by the length-dependent mechanism until they all converge to the same depolymerization rate, which has a fixed value (Fig. 2c, bottom). To obtain a change in poleward flux, minus-end perturbations which directly change the depolymerization rate must be used (Fig. 2d). These results demonstrate that plus-end perturbations have no effect on poleward flux velocity when only plus-end length-dependence is included in the model.

**Table 1.**
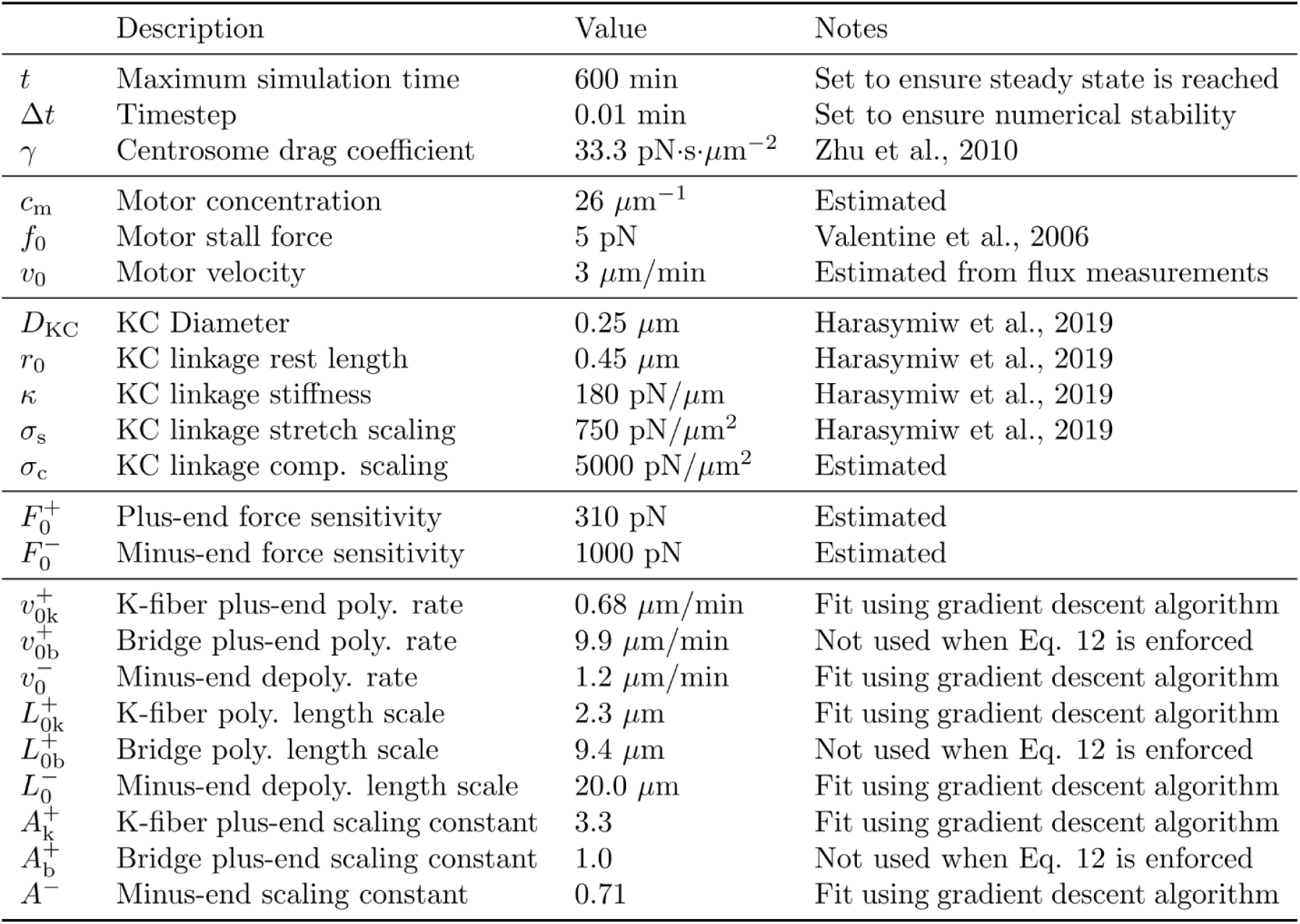
Set of parameters used in the model.

The version of the model with only plus-end length-dependence has major weaknesses when compared to previously published experiments. We see that it is sufficient to explain experiments in which minus-end perturbations, e.g., WDR62 or KATNB1 depletions, result in spindle length increasing but poleward flux velocity decreasing.^41^ However, it is unable to explain experiments in which plus-end perturbations, e.g., KIF18A or KIF4A depletions, result in both spindle length and poleward flux velocity increasing.^43^ Therefore, plus-end length-dependent mechanisms on their own are not sufficient to fully recreate observed experimental results.

Finally, we consider the full model with both plus- and minus-end length-dependent mechanisms included. We encode the minus-end length-dependent mechanism in our model as a linear increase in minus-end depolymerization rate as microtubule length increases, representing continually increasing depolymerization for longer microtubules (Methods). With this mechanism added in addition to the plus-end length-dependent mechanism described earlier, we see that the spindle once again reaches a well-defined steady-state for the three different plus-end perturbations (Fig. 2e, top). However, the polymerization rates in each case now converge to a different depolymerization rate (Fig. 2e, bottom). This leads to a direct relationship between spindle length and poleward flux for plus-end perturbations (Fig. 2f). Additionally, the inverse relationship between spindle length and poleward flux for minus-end perturbations is retained. This demonstrates that a minus-end length-dependent mechanism allows the minus-end to respond to perturbations at the plus-end, consistent with previous work.^23^ More generally, these results show that length-dependent mechanisms at both microtubule ends facilitate synchronization of plus- and minus-end dynamics by allowing them to communicate to each other through the overall length of the microtubule. Based on these results, we propose that a combination of plus- and minus-end length dependent mechanisms is necessary to explain experimental results.

## Synchronization of plus- and minus-end dynamics facilitates robust spindle length control over a wide range of flux velocities

We have shown that length-dependent mechanisms at both the plus- and minus-end of microtubules allow poleward flux velocity to react to perturbations at both ends (Fig. 2f). Notably, perturbations at either the plus- or minus-end result in poleward flux velocity either increasing or decreasing for larger spindle lengths, respectively. Here, we explore the implications of these opposite trends through simultaneous perturbations of both microtubule ends. We do this by modulating parameters in our model meant to represent the concentration of proteins that accumulate in a length-dependent manner and regulate plus- or minus-end dynamics (Fig. 3, Methods). By linking our model with protein concentrations in this way, we can better interpret experimental results and make predictions that can be directly tested.

**Figure 3.**
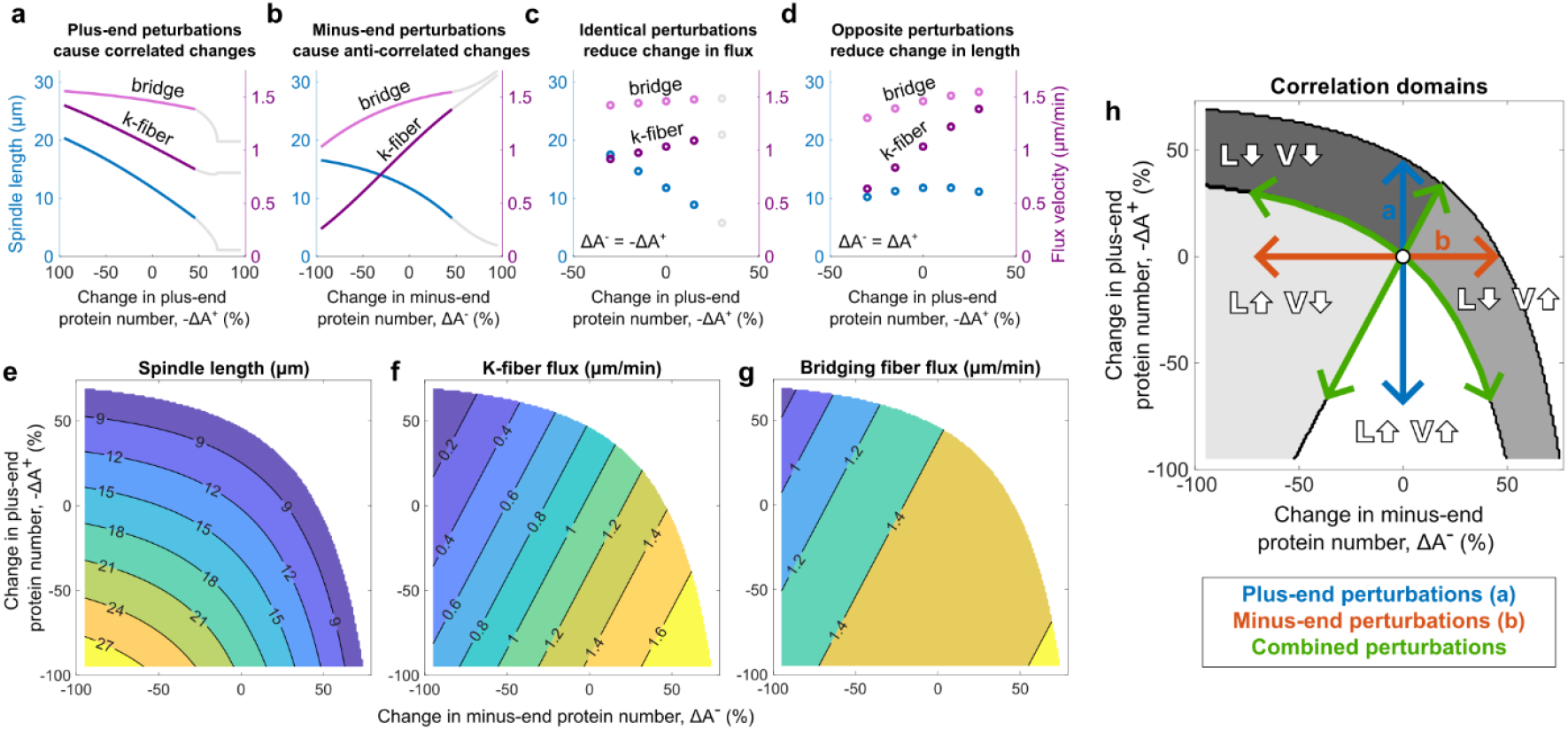
Synchronization of plus- and minus-ends through length-dependent mechanisms leads to rich dynamics in spindle length vs flux velocity. **a-d**, Plots of spindle length (blue) and poleward flux velocity (purple for k-fiber, magenta for bridging fiber) in the model for (**a**) increasing plus-protein number, (**b**) increasing minus-end protein number, (**c**) increasing both plus- and minus-end protein number by the same amount, and (**d**) increasing plus-end protein number while decreasing minus-end protein number by the same amount. Data which corresponds to spindle lengths less than 6.5 microns are shown in grey. In these plots, we assume that decreasing the parameter 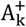 corresponds to increasing the number of plus-end-associated proteins which inhibit polymerization, e.g., KIF18A, whereas increasing A^−^ corresponds to increasing the number of minus-end-associated proteins which promote depolymerization, e.g., KIF2A, since both of these lead to smaller spindles in our model. In panels (**a**) and (**b**), the scaling constants 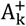 and A^−^ are varied between 5 and 195% of their nominal values (Table 1). In panels (**c**) and (**d**), the scaling constants are varied between 70% and 130% of their nominal value in discrete steps of 15%. **e-g**, Contour plots of (**e**) spindle length, (**f**) k-fiber flux velocity, and (**g**) bridging fiber flux velocity showing lines of constant value for each quantity as plus- and minus-end protein numbers are varied. Only data which corresponds to a spindle length greater than 6.5 microns are shown. **h**, Plot of correlation domains set by panels (**e**), (**f**), and (**g**). Cross sections which correspond to panels (**a**) and (**b**) appear as blue and orange lines, respectively, while lines of constant value from combined perturbations appear in green.

Consistent with our previous results, modulating the concentration of plus- and minus-end-associated proteins leads to spindle length and poleward flux having correlated and anti-correlated changes, respectively (Fig. 3a, b). Notably, the change in either quantity has a comparable magnitude for the same percent change in plus- or minus-end-associated protein number. As a result, the change in poleward flux velocity is substantially reduced if plus- and minus-end-associated protein concentrations are both varied by the same amount (Fig. 3c). Conversely, the change in spindle length is substantially reduced if plus- and minus-end-associated protein concentrations are instead varied in equal but opposite ways (Fig. 3d). The difference between plus- and minus-end perturbations thus allows for changes in either spindle length or poleward flux velocity to be partially cancelled out by combining these perturbations in different ways.

Seeing this difference in behavior motivated us to explore the full landscape of how spindle length and poleward flux velocity vary with simultaneous plus- and minus-end perturbations (Fig. 3e-g). We see that spindle length can be held fixed by compensating for an increase in protein concentration at one microtubule end with a decrease at the other (Fig. 3e). On the other hand, poleward flux velocity can be held fixed by increasing or decreasing protein concentrations at both ends simultaneously (Fig. 3f, g). Therefore, our model suggests that the synchronization of plus- and minus-end dynamics through length-dependent mechanisms allows spindle length and poleward flux to be uncoupled from one another through simultaneous perturbations, facilitating robust spindle length control over a wide range of poleward flux velocities.

The opposing response to changes in protein concentration prompted us to define four different domains relative to our control value obtained with nominal parameters: spindle length and poleward flux both increase, spindle length and poleward flux both decrease, spindle length increases while flux decreases, and spindle length decreases while flux increases (Fig. 3h). Our model predicts that overexpression or depletion of plus- or minus-end proteins each have distinct outcomes that correspond to one of the four correlation domains. We want to substantiate this prediction by testing perturbations of other parameters related to microtubule dynamics in our model as a control. Indeed, we observe the same qualitative trends if we independently vary either the strength of the length-dependent mechanisms (Fig. S1a-h) or baseline constant polymerization and depolymerization rates (Fig. S1i-p). Additionally, the same trends appear if overlap length is allowed to vary freely (Fig. S2a-h), i.e., the plus-end polymerization rate of bridging fibers is not constrained (Methods, see Eq. 12). Notably, pathways for constant spindle length and constant poleward flux velocity in addition to the four distinct correlation domains appear in all cases, indicating that this behavior occurs for generalized perturbations of microtubule dynamics and is independent of overlap length being held fixed. Therefore, the conclusions presented here do not depend on our choice of perturbations or the constraint of having a fixed overlap length.

## Length- and force-dependent mechanisms work together to make bridging fiber flux more resilient to change

In our results so far, bridging fiber flux has been more resistant to change than k-fiber flux (Fig. 3, Fig. S1 and S2). Additionally, changes in bridging fiber flux appear to be asymptotic, becoming substantially reduced as higher values are approached (Fig. 3g). This is consistent with previous experiments which showed that bridging fiber flux did not substantially increase even as k-fiber flux did.^43^ These results suggest that there is some process which regulates bridging fiber flux to make it resilient to perturbations and establishes some upper limit.

To investigate these trends in poleward flux velocity, we varied the baseline depolymerization rate in our model and observed how microtubule length and force on the minus-ends responds, since both of these quantities appear in the factors which modulate poleward flux velocity (Methods, see Eqs. 10 & 11). We see that the lengths of both microtubules decrease as the baseline depolymerization rate increases, and therefore the minus-end length-dependent factor decreases for both k-fibers and bridging fibers (Fig. 4a). Force generation from motors also decreases as depolymerization rate is increased (Methods, see Eqs. 5 & 8) and so the force-dependent factor decreases as well. However, the forces experienced by bridging and k-fiber minus-ends are in opposite directions due to the intrinsic geometry of our model, so this decrease in the force-dependent factor leads to k-fiber flux increasing but bridging fiber flux decreasing (Fig. 4b). Thus, the force- and length-dependent mechanisms partially cancel each other out for k-fibers, leading to a near-linear increase in flux velocity as baseline depolymerization rate increases (Fig. 4c). Conversely, these two mechanisms add together and substantially decrease bridging fiber flux as baseline depolymerization rate increases, leading to much smaller changes. Force- and length-dependent factors therefore work together to keep bridging fiber flux constant through perturbations of baseline depolymerization rate in our model.

**Figure 4.**
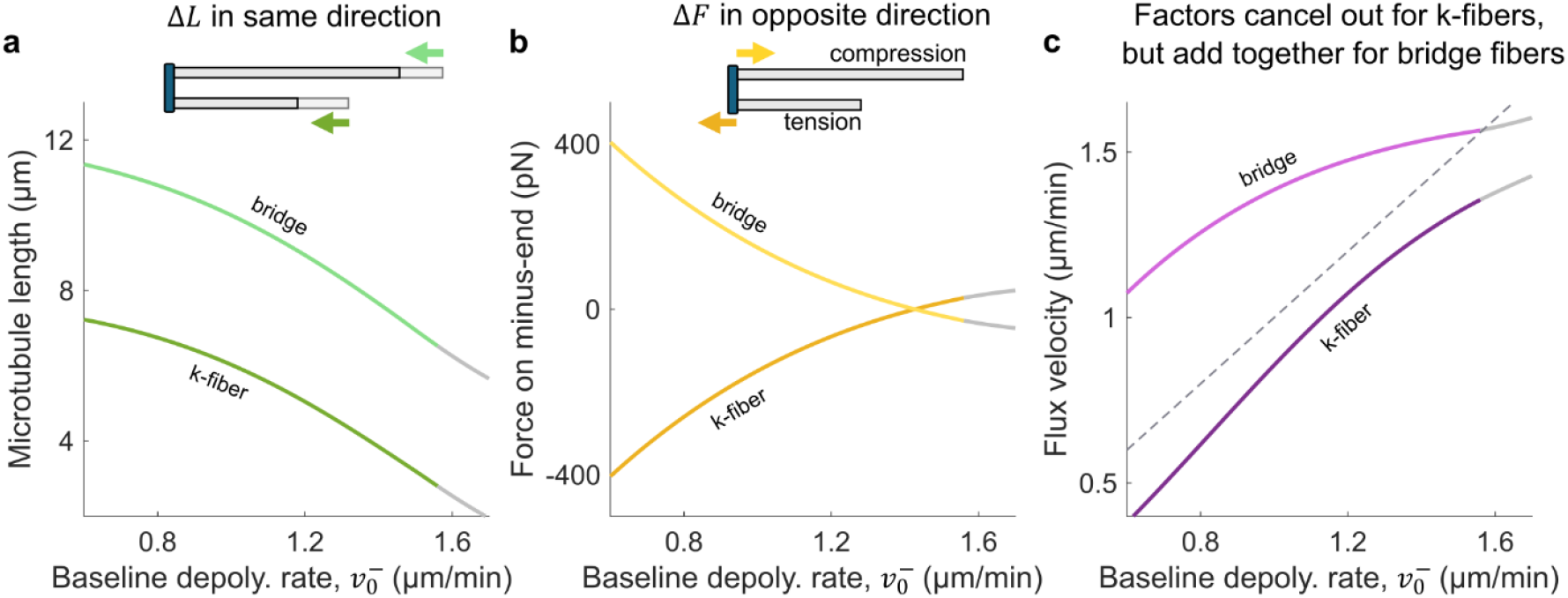
Force- and length-dependent mechanisms work together to keep bridging fiber flux constant as k-fiber flux changes. **a**, Plot of microtubule lengths for the left bridging (light green) and left k-fiber (dark green) as baseline depolymerization rate 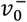 is varied. Schematic shows that the changes in length for both the bridge and k-fiber are in identical directions. **b**, Plot of force experienced by minus-ends of the left bridge (light yellow) and left k-fiber (dark yellow) as baseline depolymerization 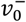 rate is varied. Schematic shows that changes in force for the left bridge and k-fiber are equal in magnitude but opposite in direction. **c**, Plot of steady-state poleward flux velocity for the left bridge (magenta) and left k-fiber (purple) as baseline depolymerization rate 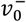 is varied. Dashed grey line corresponds to baseline depolymerization rate, i.e., has a slope of 1, which both flux velocities would be equal to if they were not modulated by length- or force-dependent mechanisms.

These results also show that, as poleward flux velocity exceeds the stepping velocity of motors, the forces produced by motors invert and begin to pull centrosomes together rather than pushing them apart. We observe this in Fig. 4b when bridging and k-fiber forces cross over zero, indicating that the direction of forces in the spindle have been reversed. Shortly after this crossover, the spindle collapses. Our model thus predicts that the upper limit of flux velocity ultimately comes from the stepping velocity of motors. A corollary of this result is that the length-dependent mechanism at the minus-end acts to modulate forces within the spindle by increasing poleward flux velocity for longer microtubules with larger overlaps. Without this modulation, increasing motor forces would decrease k-fiber flux, resulting in a feedback loop that leads to significantly increased forces (Fig. S3). Therefore, a length-dependent mechanism at microtubule minus-ends helps maintain constant forces within the spindle over a wider range of conditions.

## In-vivo depletion experiments confirm predictions of model

Our model predicts that perturbations at microtubule plus-ends lead to correlated changes in spindle length and poleward flux velocity, perturbations at minus-ends lead to anti-correlated changes, and simultaneous perturbations at both ends can allow one quantity to remain approximately constant while the other varies. In other words, our model predicts that changes in proteins concentrations influence spindle length and poleward flux in distinct ways based on whether they predominantly affect plus- or minus-end dynamics. A consequence of this is that each pair of spindle length and poleward flux values corresponds to a unique coordinate of plus- and minus-end protein concentrations (Fig. 3). Based on this, we develop a method for quantitatively comparing our model predictions against experiments which perturb proteins known to regulate microtubule dynamics. We use experimental measurements of spindle length and poleward flux as input values, and our model assigns coordinates to them that predict the extent to which plus- and minus-end dynamics are affected without any prior knowledge of biological function. This allows us to not only test our model against known protein function but also extrapolate and infer the function of proteins which are less well understood.

In order to test the predictions of our model, we utilize proteins which have well known biological functions: the plus-end polymerization inhibitor KIF18A,^56^ the minus-end severing protein KATNB1,^59^ and the minus-end depolymerizing protein KIF2A.^60^ We performed siRNA-mediated single depletion experiments *in vivo* using hTERT RPE1 cells and measured resulting spindle length and poleward flux velocity (Fig. 5a, b, Fig. S4, Fig. S5). We map these measurements onto unique coordinates in the correlation domains predicted by our model to see how the two compare (Fig. 5c, Fig. S6). Based on their locations in this plot, our theory suggests that KIF18A predominantly affects plus-end dynamics, whereas both KATNB1 and KIF2A predominantly affect minus-end dynamics. This is consistent with their known biological function, corroborating the basic assumptions of our model.

**Figure 5.**
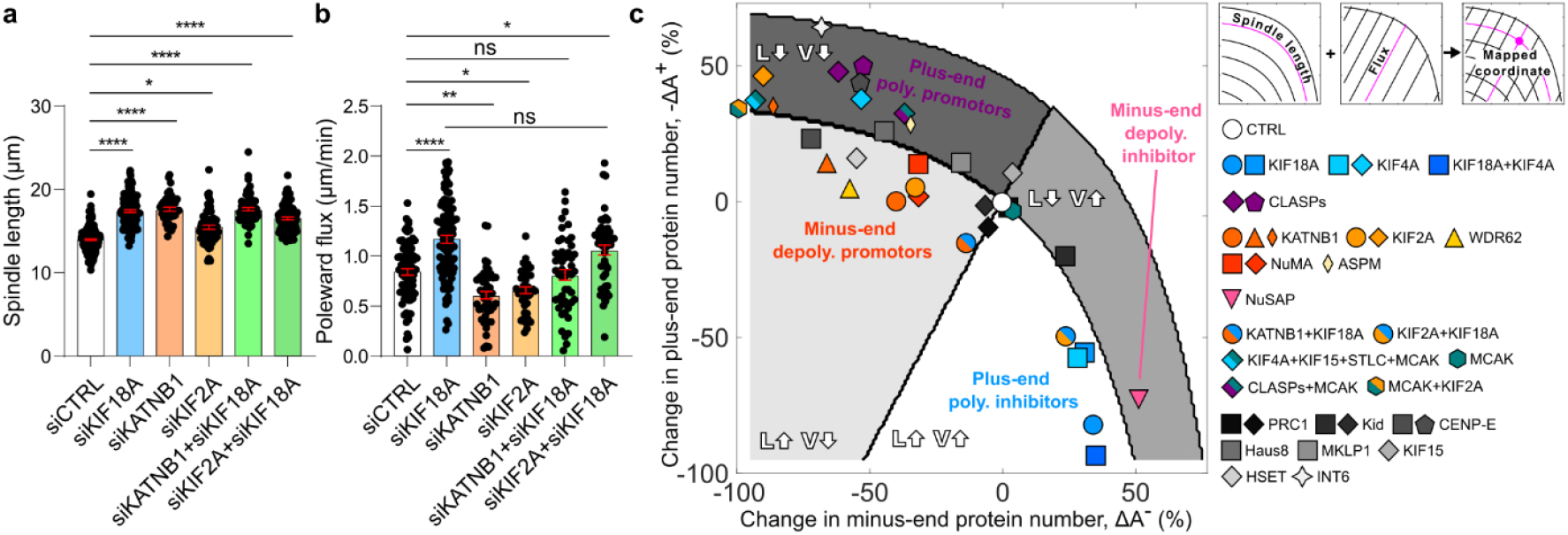
Depletion experiments confirm trends in spindle length versus flux velocity predicted by model. **a** and **b**, Experimental measurements of (**a**) spindle length and (**b**) poleward flux velocity in human cells for various siRNA-mediated proteins depletions. Mean values of spindle length are: siControl (N = 198) 14.4 ± 0.30 μm, siKIF18A (N = 133) 17.48 ± 0.25 μm, siKATNB1 (N = 42) 17.75 ± 0.48 μm, siKIF2A (N = 64) 16.52 ± 0.93 μm, siKATNB1+siKIF18A (N = 72) 17.88 ± 0.63 μm, and siKIF2A+siKIF18A (N = 90) 16.47 ± 0.30 μm. Mean values of poleward flux velocity are: siControl (N = 83) 0.84 ± 0.03 μm/min, siKIF18A (N = 96) 1.16 ± 0.05 μm/min, siKATNB1 (N = 50) 0.62 ± 0.03 μm/min, siKIF2A (N = 41) 0.65 ± 0.05 μm/min, siKATNB1+siKIF18A (N = 48) 0.80 ± 0.05 μm/min, and siKIF2A+siKIF18A (N = 51) 1.07 ± 0.02 μm/min. **c**, Data from this work (circles) and previously published experiments (other symbols) mapped onto the correlation domains predicted by our model (Fig. 3g) using overall flux values from theory (Fig. S6). Depletions of proteins known to inhibit or promote plus-end polymerization are colored in shades of blue and purple, respectively, whereas those known to inhibit or promote minus-end depolymerization are colored in shades of pink and red/orange, respectively. The protein MCAK, which affects both microtubule ends, is colored green. Co-depletions are represented by the colors of their constituent single depletions. Other MAPs which do not directly affect microtubule dynamics are depicted in shades of grey. Circles correspond to original data from panels (**a**) and (**b**), and other symbols represent data from previously published work: squares are Risteski et al., 2022; square diamonds are Steblyanko et al., 2020; thin diamonds are Jiang et al., 2017; triangles are Guerreiro et al., 2021; pentagons are Maffini et al., 2009; hexagons are Ganem et al., 2005; upside-down triangles are Sun et al., 2024; and four-pointed stars are Renda et al., 2017. Note that experimental measurements are normalized relative to theory such that the reported control value corresponds to simulation output using nominal parameter values (Table 1).

We next wanted to explore simultaneous perturbations of plus- and minus-end dynamics, which our theory predicts can allow spindle length and poleward flux to vary independently. We do this by using two siRNA-mediated co-depletion experiments (Fig. 5a, b, Fig. S4, Fig. S5). When KIF18A and KATNB1 are co-depleted, we see that spindle length increases, but remarkably no significant change in poleward flux velocity occurs. On the other hand, when KIF18A and KIF2A are co-depleted, we see that both spindle length and poleward flux velocity increase. As before, we map these measurements onto theoretically predicted coordinates (Fig. 5c, Fig. S6). The locations of these depletions in our correlation domains indicate that the KIF18A+KATNB1 co-depletion affects plus- and minus-end dynamics to a similar extent, whereas the KIF18A+KIF2A co-depletion predominantly affects plus-end dynamics. Our results therefore suggest that while KIF18A and KATNB1 have comparable contributions to microtubule dynamics, KIF18A has a stronger effect than KIF2A. All together, these results show that depletion of plus- and minus-end-associated proteins can be simultaneously combined to vary spindle length and poleward flux independently of one another *in vivo*, confirming our model predictions.

Finally, we map depletions of relevant proteins from previous experiments onto the correlation domains predicted by our model (Fig. 5c). We include experiments that measured spindle length and poleward flux velocity for different perturbations that affected spindle dynamics.^35,41–47^ Depletions which involve NDC80 are excluded since they eliminate k-fibers from the spindle^45^ and our model is unable to represent such geometry. In these experiments, the depleted proteins known to be associated with plus-end regulation are KIF18A,^56^ KIF4A,^61^ and CLASPs.^64^ Since the plus-end length-dependent mechanism in our model is based on inhibitors such as KIF18A and KIF4A, depletion of these proteins directly corresponds to a reduction of plus-end-associated proteins in the model. On the other hand, depletion of polymerization promoters such as CLASPs corresponds to an increase of plus-end-associated proteins in the model. Indeed, both trends are shown when mapping these depletion experiments onto our model. Proteins known to be associated with minus-end dynamics in these experiments are KATNB1,^59^ KIF2A,^39^ NuMA,^62^ WDR62,^63^ ASPM,^47^ and NuSAP.^46^ All of these proteins effectively increase minus-end depolymerization rate with the exception of NuSAP, which inhibits depolymerization by KIF2A in addition to other functions. Consistent with our predictions, the first five depletions are mapped onto positions which correspond to a lower concentration of minus-end depolymerizing proteins in our model, whereas NuSAP depletion corresponds to a higher concentration. Co-depletion experiments are mapped onto locations that are between the locations of their constituent single depletions, and so our model suggests that they partially cancel each other out. Depletion of proteins which affect both microtubule ends, e.g., MCAK,^64^ or do not directly influence microtubule dynamics at all, e.g., Haus8^65^ or PRC1, are difficult to interpret using our model but are still included for the sake of completeness. These results show that our model can rectify the seemingly contradictory results of how spindle length and poleward flux change relative to one another in these experiments.

We note that, for most single depletion experiments, the mapped coordinates correspond to changes in both plus- and minus-end protein concentration in our model. In almost all cases, the protein concentration most substantially changed in our model coincides with the known function of the depleted protein. We propose that the smaller change in protein concentration at the opposite end ultimately arises from simplifications used in our model, which describes complex physical processes as linear relationships. Differences between individual depletions of the same protein are attributed to differences in microtubule regulation between cell lines, e.g., RPE1 cells used in Risteski et al. whereas Steblyanko et al. uses U2OS cells. The only data point which lies outside the bounds of our model domains is the MCAK+KIF2A co-depletion from Ganem et al., which has a measured poleward flux velocity that is slightly smaller than the lower bound in our model. We attribute this to a breakdown of the assumptions in our model for exceedingly low minus-end protein concentrations, i.e., near a 100% reduction in protein number. All together, these results validate the prediction of our model that plus- and minus-end perturbations have distinct effects on how flux velocity changes with spindle length. Therefore, our model can explain the vast majority of depletion experiments across these numerous previous works.

## Discussion

We have shown that length-dependent mechanisms can synchronize the dynamics of microtubule plus- and minus-ends, resulting in them working together rather than length being regulated entirely by the plus-end.^54,56,57^ This synchronization can explain changes in poleward flux velocity resulting from depletion of plus-end-associated proteins and, in general, how the spindle regulates changes in flux and maintains a stable geometry over a wide range of flux values. Our model predicted that plus-end perturbations lead to correlated changes in spindle length and poleward flux, whereas minus-end perturbations lead to anti-correlated changes, and we validated these predictions with *in vivo* depletion experiments of the plus-end-associated protein KIF18A and minus-end-associated proteins KIF2A and KATNB1. Finally, we demonstrated that simultaneous perturbations of plus- and minus-ends can cancel out changes in either spindle length or flux velocity in our model, which we corroborated with an *in vivo* double depletion experiment that rescues poleward flux velocity for larger spindle length in human cells.

Our results provide an explanation for past experiments in which spindle length and poleward flux were observed to change in the same direction,^40,42–45^ in the opposite direction,^34,41^ or vary independently of one another.^35,47^ In this paper, we demonstrated that synchronization of both microtubule ends through length-dependent mechanisms allows these seemingly contradictory results to be rectified through perturbations that predominantly affect microtubule plus-ends, minus-ends, or a combination of both, respectively. This expands upon previous work which proposed a direct relationship between spindle length and poleward flux velocity.^23,43,45^ Additionally, our results shed light on how poleward flux can vary by an order of magnitude between species despite having similar spindle lengths.^29,31,43^ More generally, we propose that this synchronization mechanism allows cells to better maintain their spindle length across varying conditions (Fig. 6). Loss of proteins at either end would lead to a larger spindle, causing more proteins to accumulate at the opposite end and increase length regulation, reducing overall spindle size at the cost of poleward flux velocity. Additionally, cells could directly upregulate minus-end-associated proteins to compensate for a loss of plus-end-associated proteins or vice-versa, maintaining a constant spindle length in exchange for changing poleward flux velocity. Thus, our work not only resolves a paradox of how spindle length and poleward flux relate to each other by unifying different observed trends, but it also reveals a fundamental mechanism of spindle length control.

**Figure 6.**
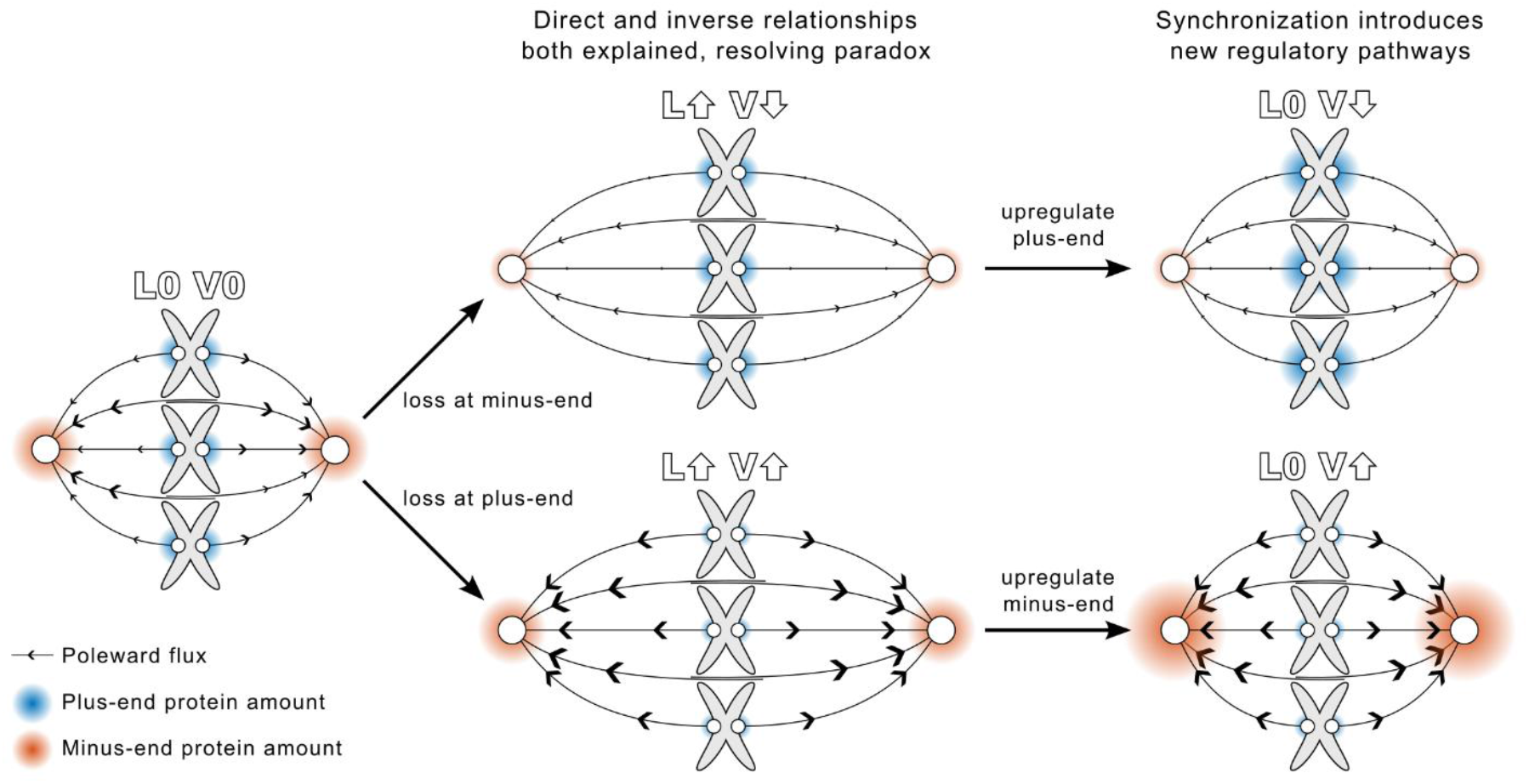
Schematic demonstrating robust spindle length control via the synchronization of microtubule plus- and minus-ends. Length-dependent mechanisms at both microtubule ends allows each end to sense and respond to changes at the opposite end. Loss of either minus- (top) or plus-end-associated (bottom) proteins leads to larger spindles and decreased or increased poleward flux velocity, respectively. The synchronization of plus- and minus-end dynamics from length-dependent mechanisms allows cells to upregulate proteins at the opposite end to exert increased length regulation, partially or fully rescuing spindle length at the cost of poleward flux velocity decreasing (top) or increasing (bottom).

The synchronization mechanism uncovered in this paper also answers a longstanding question of how microtubule ends coordinate with one another despite being separated by microns. In anaphase, for example, chromosome segregation is thought to be controlled by the coordinated action of both plus- and minus-end-associated proteins,^66– 68^ and the process is followed by a reduction of poleward flux velocity and spindle elongation across species.^33,69^ Our model shows that synchronization of plus- and minus-end dynamics through length-dependent mechanisms at both microtubule ends would facilitate this type of coordinated behavior. Furthermore, it is known that the reduction of poleward flux and subsequent spindle elongation is driven by downregulation of kinesin-13 proteins,^70,71^ which is consistent with the predictions of our model. Additionally, our model predicts that lowering poleward flux velocity allows motor proteins in antiparallel overlaps to exert significantly more force, further contributing to rapid elongation of the spindle and segregation of chromosomes. Our proposed synchronization mechanism therefore provides additional insight into how transitions in spindle length are controlled and how plus- and minus-end-associated proteins work together in a precise way. Furthermore, the conclusions presented here can be extended to other systems which exhibit flux of microtubule, e.g., developing neurons where continual retrograde flow of microtubules towards the soma is involved in axon formation.^72,73^

This work lays the groundwork for a fundamental mechanism of spindle length regulation and opens new directions for future studies. Follow-up experiments that combine depletion of one protein with overexpression of another could offer additional insight into the cellular function of the regulatory pathways proposed in this paper. Additionally, experimentally identifying the proteins involved in our hypothesized minus-end length-dependent mechanism would provide additional context for how cells utilize this mechanism. Expanding our model, such as incorporating dynamic properties of microtubule that result in different distributions, would allow us to explore the effects of our proposed synchronization mechanism in different stages of mitosis, e.g., prometaphase when k-fibers are forming and anaphase when sister kinetochores are no longer linked. Furthermore, adding dynamic kinetochore-microtubule attachments would let us investigate chromosome oscillations and how they respond to the perturbations described in this paper. Finally, incorporating additional spatial dimensions and steric interactions between chromosomes would let us study aberrant architectures such as multipolar spindles, and the mechanisms of length control proposed here may help characterize how these structures evolve over time and congregate their excess centrosomes.^74–76^

## Methods

### Model equations of motion

To describe the geometry of the system, we follow the position of the left and right centrosomes, denoted as *x*_*ℓ*_ and *x*_r_, respectively, over time *t*. We choose a coordinate system such that *x* = 0 is the center of the spindle and the left (right) region corresponds to negative (positive) positions and direction. Each term is denoted with an *ℓ* or r subscript to indicate which region it is in. The length of the spindle, *L*, is defined as the separation between centrosomes, *L* = *x*_r_ − *x*_*ℓ*_, which is strictly positive due to our choice of coordinate system. The coordinates *x*_*ℓ*_ and *x*_r_ also correspond to the position of minus-ends for microtubules originating from the left and right centrosome, respectively, since we assume that they are permanently bound together. Microtubule fibers are thus fully defined by their lengths, *L*_*j*_ for index *j* = b*ℓ*, k*ℓ*, br, kr corresponding to left bridging, left k-fiber, right bridging, and right k-fiber, respectively, which we use to calculate plus-end positions in conjunction with centrosome position, e.g., 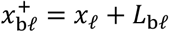 is the position of the left bridging fiber plus-end. With this information, we can calculate the length of all anti-parallel overlaps in the system and determine the sliding forces generated by motors, e.g., *O*_c_ = *L*_b*ℓ*_ + *L*_br_ − *L* is the length of the central anti-parallel overlap formed by the left and right bridging fibers. Similarly, we can use the position of k-fiber plus-ends to calculate interkinetochore stretch. The geometry of the system is thus fully defined by the two centrosome positions and four lengths of constituent microtubules, which we can use to calculate forces from sliding motors and kinetochore stretch.

We assume that the forces on each centrosome are balanced, i.e., the drag force acting on centrosomes is equal to the net force exerted by microtubules:

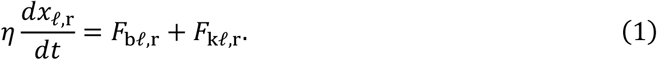

where *η* is the viscous drag coefficient, and the forces exerted on centrosomes by the bridging and k-fibers are denoted as *F*_b*ℓ*,r_ and *F*_k*ℓ*,r_, respectively. For bridging fibers, these forces come exclusively from motors in the two anti-parallel overlaps formed with opposing microtubules. For k-fibers, the force from interkinetochore stretch must also be considered. For the sake of both simplicity and clarity, we only discuss equations for the left centrosome here. The force on the left centrosome from the left bridging fiber is:

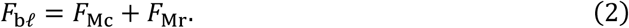

Here, *F*_Mc_ is the force generated by motors in the central overlap, and *F*_Mr_ is the force generated by motors in the right overlap. Similarly, the force exerted on the left centrosome by the left k-fiber is:

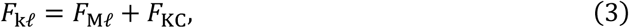

where *F*_M*ℓ*_ is the force generated by motors in the left overlap and *F*_KC_ is the tensile force between sister kinetochores. We can therefore use Eqs. (1-3) to calculate the velocity of each centrosome from the forces generated by sliding motors in anti-parallel overlaps and interkinetochore stretch.

Experiments have shown that interkinetochore forces can be characterized by an effective stiffness that depends on interkinetochore stretch.^77^ Thus, we model the linkage as a non-linear Hookean spring, giving us:

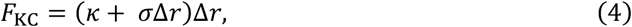

where *κ* is the stiffness of the spring, *σ* controls the nonlinear response to stretching, and Δ*r* = *r* − *r*_0_ is the stretch from rest length, *r*_0_, of the spring connecting sister kinetochores. Similar to how anti-parallel overlaps are calculated, we obtain kinetochore extension from the geometry of the system, *r* = *L* − *L*_k*ℓ*_ − *L*_kr_ − 2*D*_KC_, where *L*_k*ℓ*_ and *L*_kr_ are the lengths of the left and right k-fibers, respectively, and *D*_KC_ is the diameter of kinetochores. Here, we have assumed that the drag force of kinetochores is negligible.

To describe the force generated by motors in anti-parallel overlaps, we use a mean-field approach. First, we assume that all motors exert an equal force *f*_m_ such that *F*_M_ = *N*_m_*f*_m_, where *N*_m_ is the total number of motors in the overlap. Second, we assume that the number of motors is equal to a linear density *c*_m_ multiplied by the length of the overlap *O*, or *N*_m_ = *c*_m_*O*. Finally, we use a linear force velocity relationship, *υ*_m_ = *υ*_0_(1 − *f*_m_/*f*_0_), to describe the force generated by each motor as a function of stepping velocity *υ*_m_. Here, *υ*_0_ is the stepping velocity of motors in the absence of force and *f*_0_ is the motor stall force. Putting this all together, the motor forces on the left centrosome are:

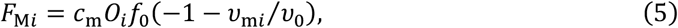

where *O*_*i*_ denotes the length of the overlap region and *υ*_m*i*_ denotes the stepping velocity of motors for index *i* = *ℓ*, c, r corresponding to left, central, and right overlap regions, respectively. The central overlap is calculated as *O*_c_ = *L*_b*ℓ*_ + *L*_br_ − *L* and the off-center overlaps are calculated as *O*_*ℓ*,r_ = *L*_k*ℓ*,r_ + *L*_br,*ℓ*_ − *L*, where *L*_b*ℓ*_, *L*_br_, *L*_k*ℓ*_, and *L*_kr_ are the lengths of the left bridging, right bridging, left k-fiber, and right k-fiber microtubules, respectively. We enforce that all overlaps are bounded between 0 and *L*, i.e., *O*_*i*_ = *max*(0, *min*(*L, O*_*i*_)). In our model, linear density *c*_m_ remains constant through time. The stepping velocity of motors can be obtained using the poleward flux velocities of microtubules, as discussed in the next section.^52^

### Model description of microtubule plus- and minus-end dynamics

To fully describe spindle length regulation, we also include dynamic instability of microtubules. We do not model the full stochastic process involving catastrophe, pause, and rescue events, but instead streamline these into an average polymerization rate at the plus-end of microtubules. We also assume that the minus-ends of microtubules are continually depolymerized by proteins at the centrosome, yielding

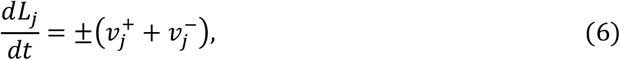

where *L*_*j*_ is the length of microtubules for index *j* = *bℓ, kℓ, br, kr* that corresponds to left bridging, left k-fiber, right bridging, and right k-fiber, respectively. Here, ± indicates a positive (negative) sign for left (right) microtubules, 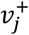 is the polymerization rate of the microtubule plus-end, and 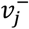 is the depolymerization rate of the microtubule minus-end. Minus-ends remain fixed to centrosomes in our model, so the length change calculated from Eq. (6) is applied entirely at the plus-end, and the sign of each individual rate is given by the direction it causes the plus-end to be displaced by. Thus, for microtubules originating from the left (right) centrosome, the depolymerization rate is negative (positive) and the polymerization rate is positive (negative). For all microtubules, regardless of which centrosome they originate from, a positive value of *dL*_*j*_/*dt* indicates an increase in length whereas a negative value indicates a decrease in length.

In our model, microtubule plus-ends move toward the centrosome they originate from if *υ*^+^ < *υ*^−^, away if *υ*^+^ > *υ*^−^, and stay at a constant distance if *υ*^+^ = *υ*^−^. Any given segment along the microtubule will move towards the centrosome with velocity *υ*^−^ regardless of what occurs at the plus-end, resulting in poleward flux (Fig. 1, inset 1). To obtain the flux velocity as measured from the lab reference frame, we also include the corresponding centrosome velocity:

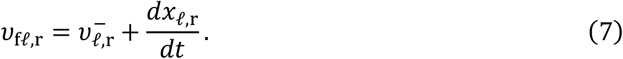

Note that, at steady state, poleward flux and depolymerization velocity are exactly equal. It can be shown that the stepping velocity of motors in an anti-parallel overlap is equal to the difference in poleward flux velocities of the microtubules which compose that overlap.^52^ For example, the motor velocity in the central overlap from the perspective of the left centrosome is given by

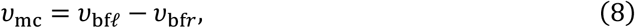

where *υ*bf*ℓ* and *υ*bf*r* are the poleward flux velocities of the left and right bridging fibers, respectively, that make up the central overlap. Note that, due to our choice of coordinate system, the flux velocity of microtubules originating from the left centrosome is always negative, whereas for microtubules originating from the right centrosome the flux velocity is always positive. If we were considering the motor velocity from the perspective of the right centrosome, the two velocities in Eq. (8) would be swapped, flipping its sign. We see that in the special case where the segment velocities as measured from the lab reference frame are equal to half of the motor stepping velocity, or *υ*_bf*ℓ*_ = −*υ*_0_/2 and *v*_bfr_ = *υ*_0_/2, the overall motor stepping velocity is −*υ*_0_ and it produces zero force as per Eq. (5). Likewise, if both poleward flux velocities are zero, the overall motor stepping velocity is zero and it produces a maximum force of −*f*_0_, which will drive the left centrosome further to the left and therefore elongate the spindle. Intuitively, this can be understood as poleward flux velocity partially canceling out the stepping velocity of motors, reducing the amount of stretch it can achieve and therefore how much force it can produce.

To investigate the physical mechanisms involved in microtubule length regulation, we also allow the baseline polymerization and depolymerization rates to be modulated by both force- and length-dependent effects. We encode these into separate factors that scale the baseline depolymerization rates as follows:

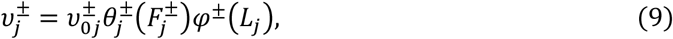

where 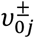 is the constant baseline polymerization and depolymerization rate of the plus- and minus-ends, respectively, and 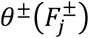 and *φ*^±^(*L*_*j*_) are functions that return a unitless scaling factor based on the force 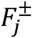 at the plus- or minus-end and length *L*_*j*_ of the microtubule, respectively.

It has been shown *in vitro* that forces affect the growing rate of microtubule plus-ends.^78^ The depolymerization of minus-ends at the centrosomes is a less well-understood topic, but it is thought that kinesin-13 proteins which localize centrosomes may ultimately be responsible for the effect.^35,79^ In this case, we assume that forces on microtubules displace the minus-ends such that they are exposed to a greater or lesser amount of kinesin-13 proteins at the centrosome, leading to a similar force dependence as plus-ends. An alternative explanation for poleward flux is that forces from sliding motors drive depolymerization at the centrosome.^43,52,70,80^ In this case, we assume a direct force dependence arises. We use a linear force-velocity relationship to encode these effects:

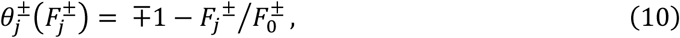

where *j* = b*ℓ*, k*ℓ*, br, kr once again corresponds to left bridging, left k-fiber, right bridging, and right k-fiber, respectively, ∓1 indicates a negative (positive) sign for microtubules originated from the left (right) centrosome, and 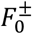 is a scaling factor that sets force sensitivity for the plus- and minus-ends, respectively. The force term *F*_*j*_^±^ corresponds to the force experienced by the plus- or minus-end, and therefore the same external force will have a different sign for either end. For example, pulling on the plus-end of the left k-fiber would result in a positive force on the plus-end and a negative force on the minus-end. A microtubule under tension would therefore experience increased polymerization at the plus-end and decreased depolymerization at the minus-end.

It is known that proteins can bind along the length of microtubules and then accumulate at the plus-end, leading to a length-dependent mechanism wherein protein number increases with microtubule length.^54^ Once microtubules reach a critical length, these proteins accumulate to a threshold level that either stalls further microtubule growth or induces catastrophe. In this manner, they establish a mechanism that sets microtubule length. Previous studies have shown that the plus-end-associated KIF18A, a protein from the kinesin-8 family, plays a dominant role in length regulation, with depletion of these proteins resulting in longer microtubules *in vivo*^55^ and longer spindles during mitosis in both fission yeasts^81–83^ and higher eukaryotes.^56,57^ The accumulation of regulating proteins on minus-ends is not as well characterized, but we hypothesize that one or more of the many proteins which localize at centrosomes could exhibit this behavior. For example, it has been shown that the abundance of both Katanin and KIF2A at a given centrosome correlates with the length of microtubules originating from it.^58^ We further propose that any protein with a long residence time can accumulate on microtubule minus-ends in a length-dependent manner due to poleward flux, consistent with previous theoretical work.^23^ Similar to how we handled force dependence, we also encode these effects into a linear function:

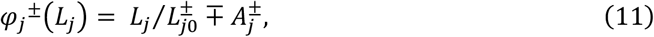

where 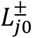 is a scaling factor that sets length sensitivity for the plus- and minus-ends, respectively, ∓ indicates a negative (positive) sign for plus-ends (minus-ends), and 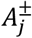 is a unitless scaling constant. For plus-ends, this function yields a positive value until 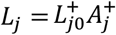, at which point it becomes negative, representing depolymerization outweighing polymerization at the plus-end for very long microtubules. For minus-ends, this function is always positive, representing a continually increasing depolymerization rate for longer microtubules. We assume that bridging and k-fibers can have different parameters at the plus-end due to interactions with kinetochores but that they have identical parameters at the minus-end since they interact with centrosomes in the same way in our model.

Overlap regulation within the mitotic spindle is currently not well understood, and therefore we choose to fix the central overlap length in our model so that effects from changing force do not influence our results. This is achieved by setting the change in length of bridging fibers equal to the velocity of the centrosome they are attached to, or

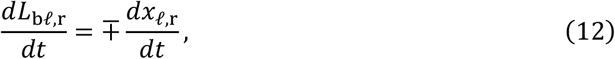

which keeps the plus-ends of bridging fibers fixed in place and therefore maintains overlap length. This allows us to study length regulation of the spindle independently of how antiparallel overlaps are maintained. However, the main conclusions of this paper are unchanged if overlap length is not fixed and instead allowed to vary freely (Fig. S2).

### Cell culture and transfection

hTERT-RPE1 cells and hTERT-RPE1 PAGFP α-Tubulin cells^84^ were cultured in Dulbecco’s Modified Eagle Medium (DMEM; ThermoFisher, 41965-039) supplemented with 10% fetal calf serum (FCS; Labforce, S181T), 100 U/mL Penicillin and 100 mg/mL Streptomycin (ThermoFisher, 15140-122) at 37°C with 5% CO2 in a humidified incubator. For live cell imaging, DMEM was replaced to Leibovitz’s L-15 medium without phenol red (ThermoFisher, 21083-027) supplemented with 10% FCS.

hTERT-RPE1 cells were transfected for 48 hours with 20nM of indicated siRNAs using Opti-MEM (ThermoFisher, 31985-062) and Lipofectamine RNAiMAX (ThermoFisher, 13778030) according to the manufacturer’s instructions. The following sense strands of validated siRNA duplexes were used: KIF2A^85^ (Ambion, 5′- GUUGUUUACUUUCCACGAATT-3′), KIF18A^56^ (Dharmacon, 5′- CCAACAACAGUGCCAUAATT-3′), KATNB1^41^ (Qiagen, 5′- GCCUGGAUUUCCACCCGUATT-3′), and as a negative control (AllStars Negative Control siRNA, Qiagen, 1027281, Proprietary). The depletion efficiency of all siRNA treatments were validated using immunofluorescence and western blotting (Fig. S5).

### Immunofluorescence

To stain for KIF2A, KATNB1, and α-Tubulin, hTERT-RPE1 cells were fixed at -20°C for 10 minutes with ice-cold methanol stored at -20°C. To stain for KIF18A and CREST, hTERT- RPE1 cells were fixed at room temperature for 15 minutes with an aqueous solution containing 4% Formaldehyde (Applichem, A3592), 20mM PIPES (Applichem, A1079.0500) pH 6.8, 10mM EGTA (Applichem, A0878.100), 0.2%-Triton X-100 (Applichem, A4975.0100), and 1mM MgCl_2_. Following fixation, cells were rinsed with PBS and blocked at room temperature for 1 hour in blocking buffer (3% BSA (Labforce, S181T) and 1% NaN_3_ (Applichem, A1430) in PBS). Cells were then incubated for 1 hour with primary antibodies diluted in blocking buffer, rinsed with PBS, and incubated for 1 hour with secondary antibodies diluted in blocking buffer. The coverslips were mounted on microscope slides (Epredia, AA00000112E01MNZ10) using Vectashield mounting medium containing DAPI (Vector Laboratories, H1200).

The following primary antibodies were used: recombinant human anti-α-Tubulin (1:500)^86^, rabbit anti-KIF18A (1:1000; Bethyl, A301-080A), rabbit anti-KIF2A (1:1000; Invitrogen, PA3-16833), rabbit anti-KATNB1 (1:250; Proteintech, 14969-1-AP), and human anti-CREST (1:500; Antibodies Incorporated). Appropriate cross-adsorbed Alexa-Fluor conjugated secondary antibodies (1:1000; Invitrogen) were used. Images were acquired using 60× (NA 1.4) oil objective on an Olympus DeltaVision wide-field microscope (GE Healthcare) equipped with a DAPI/FITC/TRITC/Cy5 filter set (Chroma Technology Corp.) and Coolsnap HQ2 CCD camera (Roper Scientific) running Softworx (GE Healthcare). Z-stacks of 12.80 µm thickness were imaged with z-slices separated by 0.2 µm. 3D images stacks were deconvolved using Softworx (GE Healthcare) in conservative mode.

### Poleward microtubule flux measurements

hTERT-RPE1 PAGFP α-Tubulin cells were cultured in 4-well glass-bottom µ-slides (Ibidi, Vitaris) and treated with siRNAs as described above. Prior to imaging, cells were incubated with 50nM SiR-DNA (Spirochrome, SC007) and 10µM MG132 for 2hr and 30min respectively. The images were acquired using the 60x (N.A. 1.4) CFI Plan Apochromat oil objective on a Nikon A1r point scanning confocal microscope equipped with a 37°C heating chamber and running NIS elements software. A rectangular ROI (250px-300px long and 5px wide) of the spindle parallel to the metaphase plate was photoactivated using a laser of wavelength 405nm at 50-100% intensity, depending on the GFP expression levels. Single focal planes were imaged every 20s for 4min. The subsequent movement of the photoactivated fluorescence towards the spindle pole was tracked over 2min, and the rate of displacement was quantified using the metaphase plate as reference point to determine the velocity of poleward microtubule flux.

### Spindle Length Measurements in Live Cells

For live cell imaging, hTERT-RPE1 cells were cultured in glass-bottom µ-slides of 2, 4, or 8 wells (Ibidi, Vitaris) and treated with siRNAs as described above. The cells were incubated with 50nM SiR-DNA (Spirochrome, SC007) and 0.5x SpY555-tubulin probe (Spirochrome, SC203) for 2hrs prior to imaging. The images were recorded using either a 40x (N.A. 0.75) or a 60x (N.A. 1.4) objective of Nikon Eclipse Ti-E wide-field microscope equipped with a GFP/mCherry/Cy5 filter set (Chroma Technology Corp.), an Orca Flash 4.0 complementary metal-oxide-semiconductor camera (Hamamatsu), and a 37°C heating chamber using NIS software. Spindle length was quantified on Imaris 7.4.2 from the time-lapse series as the distance between the two spindle poles, visualized by fluorescently labeled tubulin, in the frame immediately preceding anaphase onset.

### Statistical Tests and Analysis

All data were plotted as mean ± SEM. Appropriate statistical tests were performed and plotted in GraphPad Prism 10.6.1.

## Acknowledgements

We thank Alexey Khodjakov, Valentina Štimac, Max Schelski, and members of the Frank Bradke lab, Nenad Pavin, and Iva Tolić labs for their constructive comments on this work, as well as Ivana Šarić for assistance with assembling figures. This work was funded by the European Research Council (ERC-SyG 855158, N.P. and I.M.T.), the European Union’s Horizon Europe research and innovation programme under the Marie Skłodowska-Curie Actions (MSCA 101151485, S.A.F.), the Croatian Science Foundation (HRZZ) through Swiss-Croatian Bilateral Projects (IPCH-2022-10-9344, N.P., I.M.T., and P.M.), the Swiss National Science Foundation through a Project Grant (No. 208052, P.M.) and a Weave Grant (No. 215124, P.M. and I.M.T.), and the University of Geneva. Co-funding was provided by the Croatian Science Foundation Projects (HRZZ IP-2019-04-5967, N.P. and IP-2024-05-5336, I.M.T.) and the Croatian Government and the European Union through the European Regional Development Fund—the QuantiXLie Center of Excellence (KK.01.1.1.01.0004, N.P.). Elena Doria was supported by an EMBO short-term fellowship to visit the Tolić laboratory.

## Contributions

N.P. and P.M designed and supervised this study. S.A.F. and N.P. developed the theory. S.A.F. developed the code and performed theoretical research. S.C. and E.D. performed experiments and analyzed experimental data. S.A.F. and S.C. assembled figures. N.P., P.M. and I.M.T. contributed with ideas and discussions. The manuscript was written by S.A.F. with feedback from N.P., P.M, and I.M.T.

## Data availability

All data are provided within the article and its Extended Data files.

## Code availability

All code is available upon reasonable request.

---

**Figure S1.**
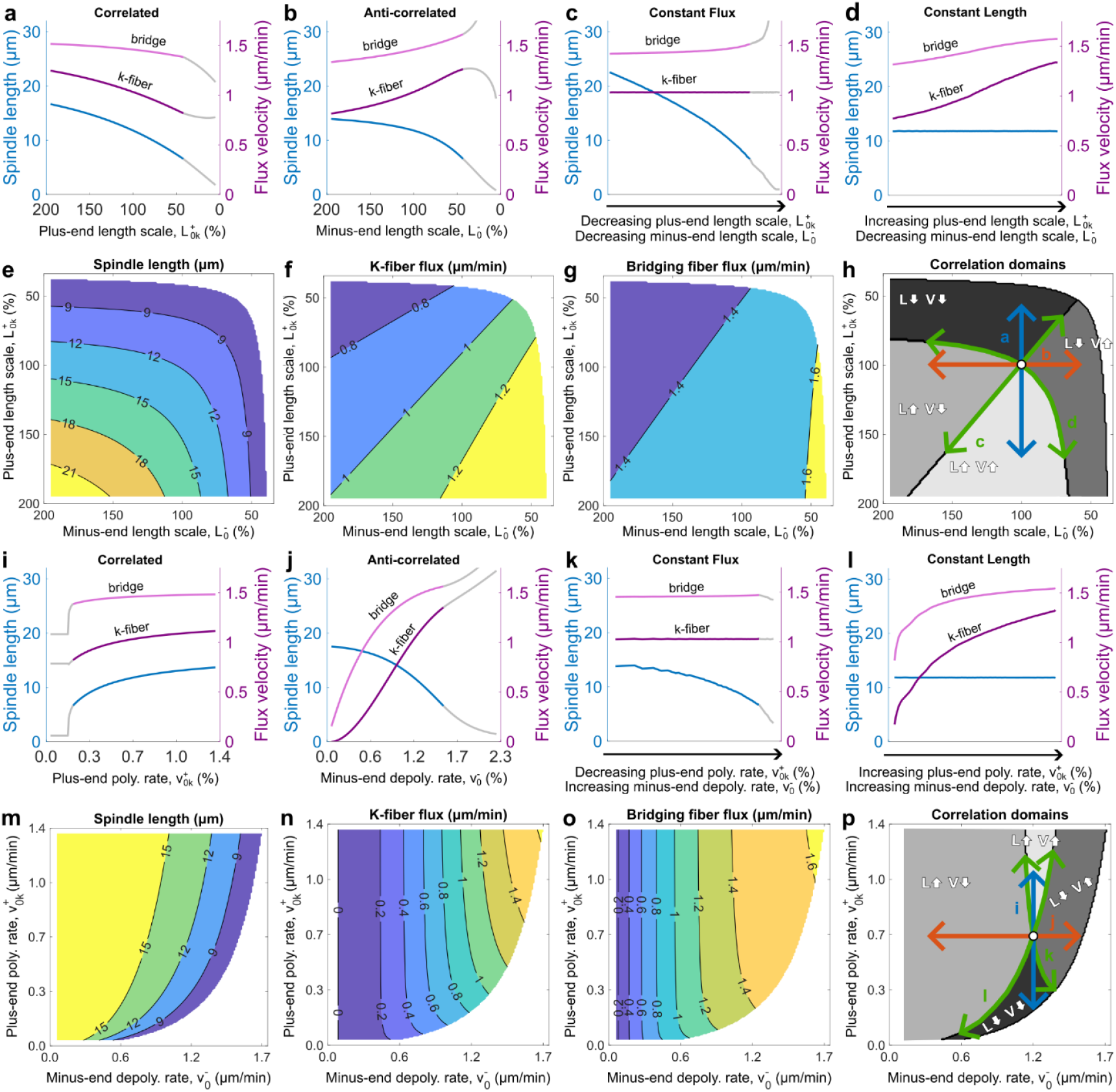
Trends in spindle length and poleward flux persist when other parameters related to microtubule dynamics are perturbed. **a-p**, Plots analogous to Fig. 3, but for length-dependence strength, 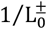 (**a-h**) and baseline polymerization and depolymerization rates, 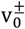 (**i-p**). In each case, lines of constant values for both spindle length and poleward flux velocity in addition to four correlation domains appear.

**Figure S2.**
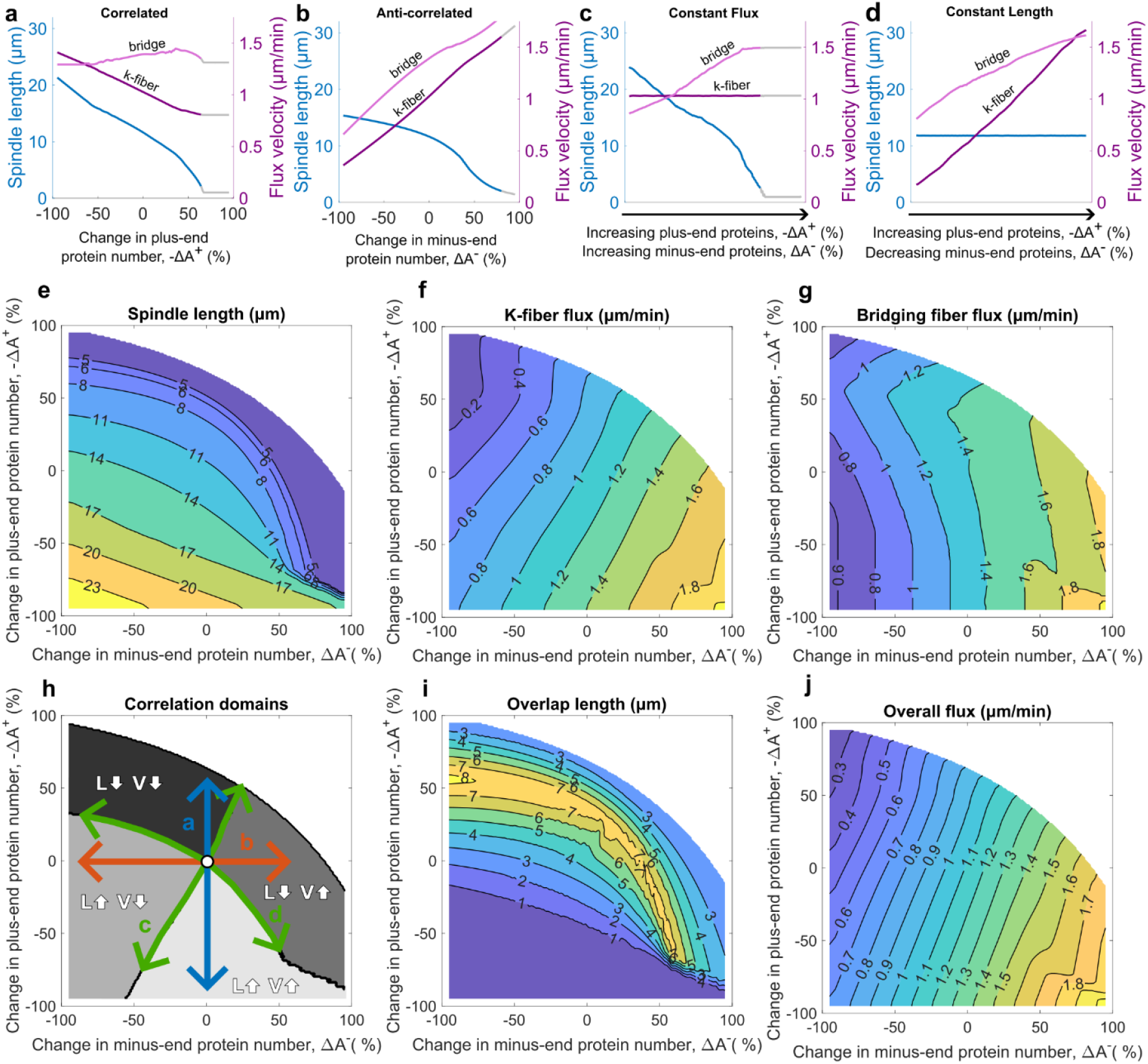
Trends in spindle length and poleward flux persist when central overlap length can vary freely. **a-h**, Plots analogous to Fig. 3, but for an overlap length that is unfixed, i.e., Eq. (12) is not enforced. We see that lines of constant values for both spindle length and poleward flux velocity in addition to four correlation domains still appear. Due to the changing overlap length, bridging fiber and k-fiber flux lines of constant value no longer have identical direction. Panel (**h**) is therefore generated using spindle length and k-fiber flux. **i**, plot of overlap length as plus- and minus-end protein number is varied. **j**, plot of overall flux as plus- and minus-end protein number is varied. Overall flux has approximately linear lines of constant value that is similar to the model with fixed overlap length. To help better visualize trends, data displayed in panels (**e-j**) has been smoothed with a window size of 0.1%.

**Figure S3.**
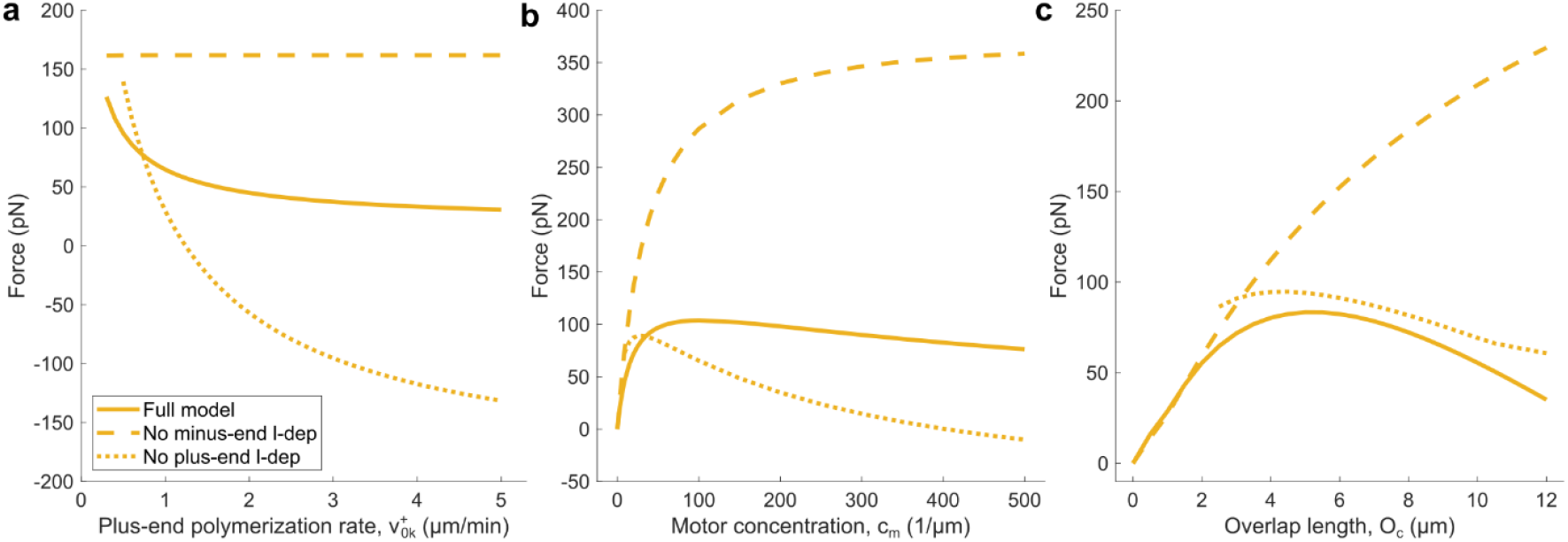
Minus-end length-dependent mechanisms work to keep forces moderate within the spindle. Force experienced by a single microtubule for varying (A) baseline polymerization rate, (B) motor concentration, and (C) overlap length. Solid lines correspond to the full model with both length-dependent mechanisms included, dashed lines correspond to the model with minus-end length dependence removed, and dotted lines correspond to the model with plus-end length-dependence removed. In all cases, removing minus-end length-dependence leads to substantially larger forces.

**Figure S4.**
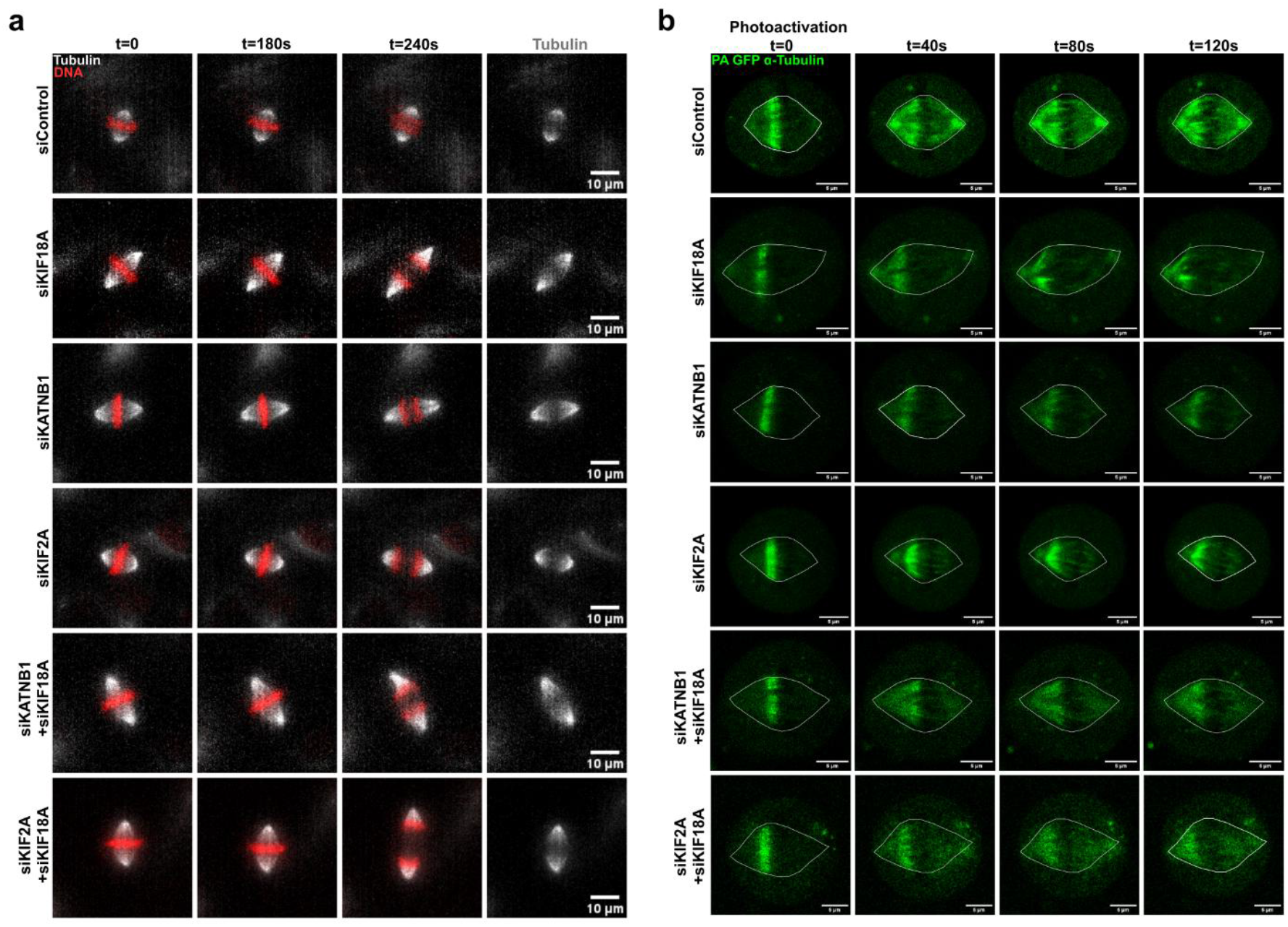
Poleward flux for different protein depletions in our experiments. **a**, Time-lapse images (maximum intensity projections) over 6 minutes of hTERT RPE1 cells treated with indicated siRNAs stained with SpY-555 Tubulin (gray) and SiR-DNA (red). All scale bars = 10 µm. **b**, Time-lapse single-plane images of photoactivated hTERT RPE1 PA-GFP α-Tubulin cells treated with indicated siRNAs over 2 minutes. All scale bars = 5 µm.

**Figure S5.**
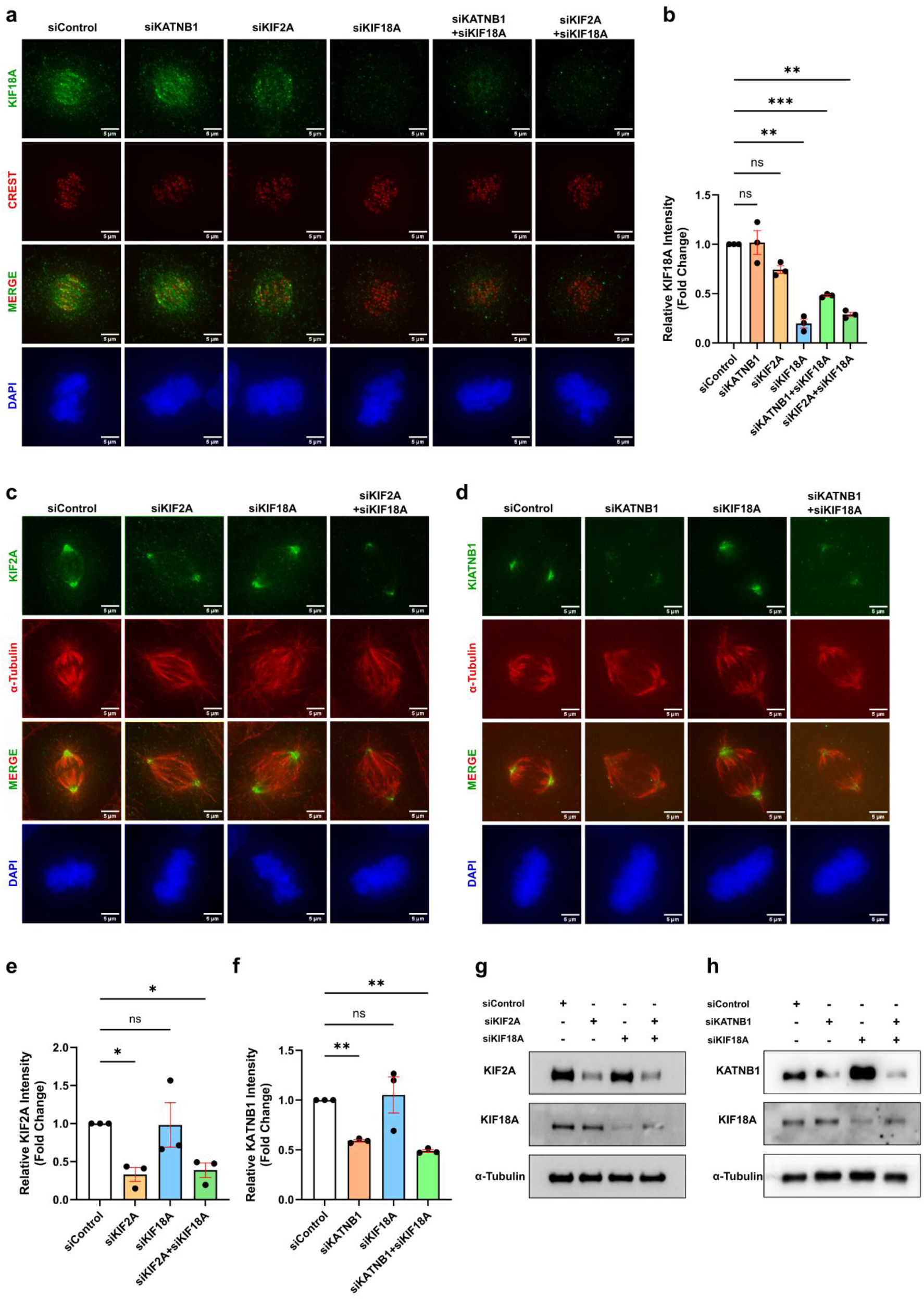
Efficiency of siRNA treatments used in our experiments. **a**, Immunofluorescence of hTERT RPE-1 cells treated with siControl, siKATNB1, siKIF2A, siKIF18A, siKATNB1+siKIF18A, siKIF2A+siKIF18A stained for KIF18A, CREST and DAPI. All scale bars are 5 µm. **b**, Quantification of normalized KIF18A intensity per cell to CREST intensity and per biological replicate to the corresponding siControl from (**a**). N = 3, n = 58-60 cells, siControl vs siKATNB1 p = 0.9998, siControl vs siKIF2A p = 0.0550, siControl vs siKIF18A p = 0.0075, siControl vs siKATNB1+siKIF18A p = 0.0009, siControl vs siKIF2A+siKIF18A p = 0.0024 in RM one-way ANOVA test. Data is represented as mean ± SEM. **c**, Immunofluorescence of hTERT RPE-1 cells treated with siControl, siKATNB1, siKIF18A, siKATNB1+siKIF18A stained for KATNB1, α- Tubulin and DAPI. All scale bars = 5 µm. **d**, Immunofluorescence of hTERT RPE-1 cells treated with siControl, siKIF2A, siKIF18A, siKIF2A+siKIF18A stained for KIF2A, α-Tubulin and DAPI. All scale bars are 5 µm. **e**, Quantification of normalized KATNB1 intensity per cell to α-Tubulin intensity and per biological replicate to the corresponding siControl from (**c**). N = 3, n = 57-63 cells, siControl vs siKATNB1 p = 0.0020, siControl vs siKIF18A p = 0.9809, siControl vs siKATNB1+siKIF18A p = 0.0015, in RM one-way ANOVA test. Data is represented as mean ± SEM. **f**, Quantification of normalized KIF2A intensity per cell to α- Tubulin intensity and per biological replicate to the corresponding siControl from (**d**). N = 3, n = 59-64 cells, siControl vs siKIF2A p = 0.0352, siControl vs siKIF18A p = 0.9993, siControl vs siKIF2A+siKIF18A p = 0.0496, in RM one-way ANOVA test. Data is represented as mean ± SEM. **g**, Western blot of hTERT RPE-1 cells treated with siControl, siKIF2A, siKIF18A, siKIF2A+siKIF18A with α-Tubulin as a loading control. **h**, Western blot of hTERT RPE-1 cells treated with siControl, siKATNB1, siKIF18A, siKATNB1+siKIF18A with α-Tubulin as a loading control.

**Figure S6.**
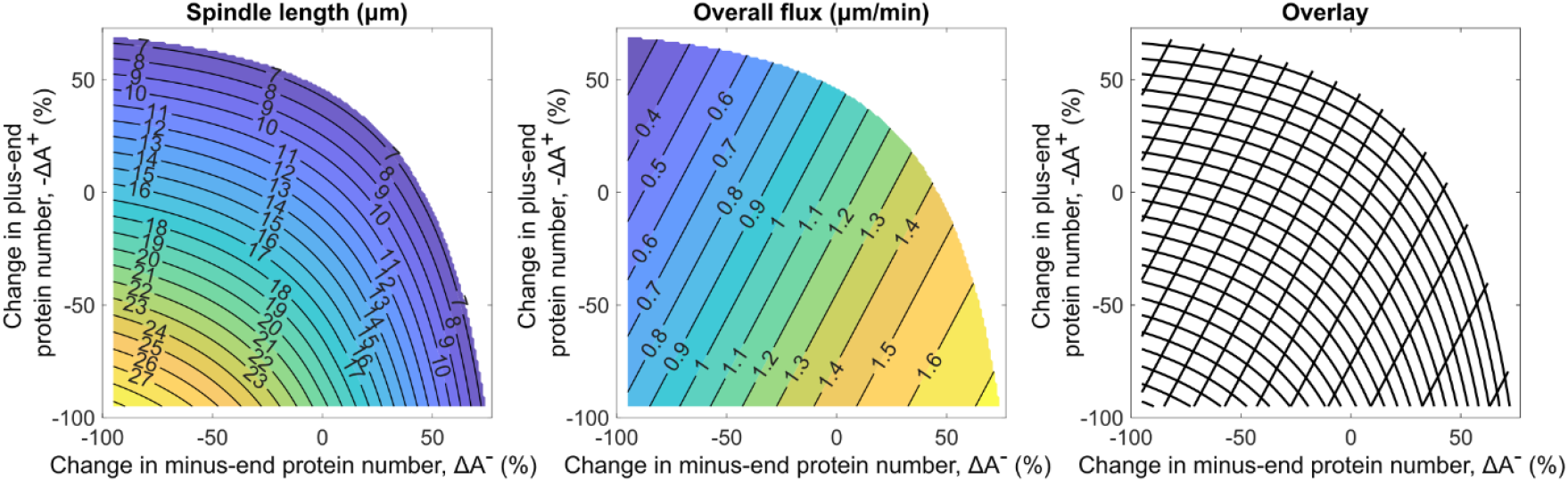
The intersection of spindle length and poleward flux lines of constant value produces a grid for quantitatively mapping experimental data. Spindle length plot is the same as Fig. 3e but with more fine grain labels. Overall flux is calculated by taking the weighted average of k-fiber flux (Fig. 3f) and bridging fiber flux (Fig. 3g) based on measurements reported in Štimac et al., 2022. A weight of 3 for k-fiber flux and 1 for bridging fiber flux was used. The overlay of these two plots produces a coordinate system in which each pair of spindle length and overall flux measurements corresponds to a unique position, allowing us to quantitatively map experimental measurements onto the predictions of our model.

## References

1. Alberts, B. et al. Molecular Biology of the Cell: Seventh International Student Edition with Registration Card. (W.W. Norton & Company, 2022).

2. Nicklas, R. B. & Staehly, C. A. Chromosome micromanipulation. Chromosoma 21, 1–16 (1967).

3. Nicklas, R. B. Measurements of the force produced by the mitotic spindle in anaphase. J Cell Biol 97, 542–548 (1983).

4. Loughlin, R., Wilbur, J. D., McNally, F. J., Nédélec, F. J. & Heald, R. Katanin Contributes to Interspecies Spindle Length Scaling in Xenopus. Cell 147, 1397–1407 (2011).

5. Lacroix, B. et al. Microtubule Dynamics Scale with Cell Size to Set Spindle Length and Assembly Timing. Developmental Cell 45, 496–511.e6 (2018).

6. Rieckhoff, E. M. et al. Spindle Scaling Is Governed by Cell Boundary Regulation of Microtubule Nucleation. Current Biology 30, 4973–4983.e10 (2020).

7. Goshima, G., Wollman, R., Stuurman, N., Scholey, J. M. & Vale, R. D. Length Control of the Metaphase Spindle. Current Biology 15, 1979–1988 (2005).

8. Wühr, M. et al. Evidence for an Upper Limit to Mitotic Spindle Length. Current Biology 18, 1256–1261 (2008).

9. Dumont, S. & Mitchison, T. J. Force and Length in the Mitotic Spindle. Current Biology 19, R749–R761 (2009).

10. Greenan, G. et al. Centrosome Size Sets Mitotic Spindle Length in Caenorhabditis elegans Embryos. Current Biology 20, 353–358 (2010).

11. Good, M. C., Vahey, M. D., Skandarajah, A., Fletcher, D. A. & Heald, R. Cytoplasmic Volume Modulates Spindle Size During Embryogenesis. Science 342, 856–860 (2013).

12. Hazel, J. et al. Changes in Cytoplasmic Volume are Sufficient to Drive Spindle Scaling. Science 342, 853–856 (2013).

13. Wilbur, J. D. & Heald, R. Mitotic spindle scaling during Xenopus development by kif2a and importin α. eLife 2, e00290 (2013).

14. Maiato, H., Gomes, A., Sousa, F. & Barisic, M. Mechanisms of Chromosome Congression during Mitosis. Biology 6, 13 (2017).

15. Decker, F., Oriola, D., Dalton, B. & Brugués, J. Autocatalytic microtubule nucleation determines the size and mass of Xenopus laevis egg extract spindles. eLife 7, e31149 (2018).

16. Valdez, V. A., Neahring, L., Petry, S. & Dumont, S. Mechanisms underlying spindle assembly and robustness. Nat Rev Mol Cell Biol 24, 523–542 (2023).

17. Zhou, C. Y. et al. Mitotic chromosomes scale to nuclear-cytoplasmic ratio and cell size in Xenopus. eLife 12, e84360 (2023).

18. Richter, M. et al. Kinetochore-fiber lengths are maintained locally but coordinated globally by poles in the mammalian spindle. eLife 12, e85208 (2023).

19. Fantes, P. A., Grant, W. D., Pritchard, R. H., Sudbery, P. E. & Wheals, A. E. The regulation of cell size and the control of mitosis. Journal of Theoretical Biology 50, 213–244 (1975).

20. Reber, S. B. et al. XMAP215 activity sets spindle length by controlling the total mass of spindle microtubules. Nat Cell Biol 15, 1116–1122 (2013).

21. Blackwell, R. et al. Physical Determinants of Bipolar Mitotic Spindle Assembly and Stability in Fission Yeast. Biophysical Journal 112, 432a (2017).

22. Oriola, D., Needleman, D. J. & Brugués, J. The Physics of the Metaphase Spindle. Annu. Rev. Biophys. 47, 655–673 (2018).

23. Wang, Y., Liu, Y.-R., Wang, P.-Y., Li, H. & Xie, P. Mechanism of spindle stability and poleward flux regulating spindle length during the metaphase. iScience 28, (2025).

24. Rubinstein, B., Larripa, K., Sommi, P. & Mogilner, A. The elasticity of motor–microtubule bundles and shape of the mitotic spindle. Phys. Biol. 6, 016005 (2009).

25. Novak, M. et al. The mitotic spindle is chiral due to torques within microtubule bundles. Nat Commun 9, 3571 (2018).

26. Margolis, R. L., Wilson, L. & Keifer, B. I. Mitotic mechanism based on intrinsic microtubule behaviour. Nature 272, 450–452 (1978).

27. Margolis, R. L. & Wilson, L. Microtubule treadmills--possible molecular machinery. Nature 293, 705–711 (1981).

28. Mitchison, T., Evans, L., Schulze, E. & Kirschner, M. Sites of microtubule assembly and disassembly in the mitotic spindle. Cell 45, 515–527 (1986).

29. Mitchison, T. J. Polewards microtubule flux in the mitotic spindle: evidence from photoactivation of fluorescence. The Journal of cell biology 109, 637–652 (1989).

30. Mitchison, T. J. & Salmon, E. D. Poleward kinetochore fiber movement occurs during both metaphase and anaphase-A in newt lung cell mitosis. J Cell Biol 119, 569–582 (1992).

31. Maddox, P., Desai, A., Oegema, K., Mitchison, T. J. & Salmon, E. D. Poleward Microtubule Flux Is a Major Component of Spindle Dynamics and Anaphase A in Mitotic Drosophila Embryos. Current Biology 12, 1670–1674 (2002).

32. LaFountain, J. R., Cohan, C. S., Siegel, A. J. & LaFountain, D. J. Direct visualization of microtubule flux during metaphase and anaphase in crane-fly spermatocytes. Mol Biol Cell 15, 5724–5732 (2004).

33. Rogers, G. C., Rogers, S. L. & Sharp, D. J. Spindle microtubules in flux. J Cell Sci 118, 1105–1116 (2005).

34. Rogers, G. C. et al. Two mitotic kinesins cooperate to drive sister chromatid separation during anaphase. Nature 427, 364–370 (2004).

35. Ganem, N. J., Upton, K. & Compton, D. A. Efficient Mitosis in Human Cells Lacking Poleward Microtubule Flux. Current Biology 15, 1827–1832 (2005).

36. Matos, I. et al. Synchronizing chromosome segregation by flux-dependent force equalization at kinetochores. J Cell Biol 186, 11–26 (2009).

37. Risteski, P. et al. Microtubule poleward flux as a target for modifying chromosome segregation errors. Proceedings of the National Academy of Sciences 121, e2405015121 (2024).

38. Maddox, P., Straight, A., Coughlin, P., Mitchison, T. J. & Salmon, E. D. Direct observation of microtubule dynamics at kinetochores in Xenopus extract spindles. The Journal of Cell Biology 162, 377–382 (2003).

39. Gaetz, J. & Kapoor, T. M. Dynein/dynactin regulate metaphase spindle length by targeting depolymerizing activities to spindle poles. The Journal of Cell Biology 166, 465–471 (2004).

40. Fu, J. et al. TPX2 phosphorylation maintains metaphase spindle length by regulating microtubule flux. J Cell Biol 210, 373–383 (2015).

41. Guerreiro, A. et al. WDR62 localizes katanin at spindle poles to ensure synchronous chromosome segregation. Journal of Cell Biology 220, e202007171 (2021).

42. Renda, F. et al. The Drosophila orthologue of the INT6 onco-protein regulates mitotic microtubule growth and kinetochore structure. PLoS Genet 13, e1006784 (2017).

43. Risteski, P. et al. Length-dependent poleward flux of sister kinetochore fibers promotes chromosome alignment. Cell Reports 40, 111169 (2022).

44. Maffini, S. et al. Motor-Independent Targeting of CLASPs to Kinetochores by CENP-E Promotes Microtubule Turnover and Poleward Flux. Current Biology 19, 1566–1572 (2009).

45. Steblyanko, Y. et al. Microtubule poleward flux in human cells is driven by the coordinated action of four kinesins. The EMBO Journal 39, e105432 (2020).

46. Sun, M. et al. NuSAP regulates microtubule flux and Kif2A localization to ensure accurate chromosome congression. Journal of Cell Biology 223, e202108070 (2024).

47. Jiang, K. et al. Microtubule minus-end regulation at spindle poles by an ASPM–katanin complex. Nat Cell Biol 19, 480–492 (2017).

48. Civelekoglu-Scholey, G., Sharp, D. J., Mogilner, A. & Scholey, J. M. Model of Chromosome Motility in Drosophila Embryos: Adaptation of a General Mechanism for Rapid Mitosis. Biophysical Journal 90, 3966–3982 (2006).

49. Civelekoglu-Scholey, G. et al. Dynamic bonds and polar ejection force distribution explain kinetochore oscillations in PtK1 cells. J Cell Biol 201, 577–593 (2013).

50. Banigan, E. J. et al. Minimal model for collective kinetochore–microtubule dynamics. Proceedings of the National Academy of Sciences 112, 12699–12704 (2015).

51. Schwietert, F. & Kierfeld, J. Bistability and oscillations in cooperative microtubule and kinetochore dynamics in the mitotic spindle. New J. Phys. 22, 053008 (2020).

52. Sigmund, I., Božan, D., Šarić, I. & Pavin, N. Mechanisms of Chromosome Positioning During Mitosis. PRX Life 2, 043017 (2024).

53. Kajtez, J. et al. Overlap microtubules link sister k-fibres and balance the forces on bi-oriented kinetochores. Nature Communications 7, 10298 (2016).

54. Howard, J. & Hyman, A. A. Microtubule polymerases and depolymerases. Curr Opin Cell Biol 19, 31–35 (2007).

55. Varga, V. et al. Yeast kinesin-8 depolymerizes microtubules in a length-dependent manner. Nat Cell Biol 8, 957–962 (2006).

56. Mayr, M. I. et al. The Human Kinesin Kif18A Is a Motile Microtubule Depolymerase Essential for Chromosome Congression. Current Biology 17, 488–498 (2007).

57. Stumpff, J., von Dassow, G., Wagenbach, M., Asbury, C. & Wordeman, L. The kinesin-8 motor Kif18A suppresses kinetochore movements to control mitotic chromosome alignment. Dev Cell 14, 252–262 (2008).

58. Thomas, A. & Meraldi, P. Centrosome age breaks spindle size symmetry even in cells thought to divide symmetrically. J Cell Biol 223, e202311153 (2024).

59. McNally, F. J. & Vale, R. D. Identification of katanin, an ATPase that severs and disassembles stable microtubules. Cell 75, 419–429 (1993).

60. Henkin, G., Brito, C., Thomas, C. & Surrey, T. The minus-end depolymerase KIF2A drives flux-like treadmilling of γTuRC-uncapped microtubules. J Cell Biol 222, e202304020 (2023).

61. Hu, C.-K., Coughlin, M., Field, C. M. & Mitchison, T. J. KIF4 Regulates Midzone Length during Cytokinesis. Current Biology 21, 815–824 (2011).

62. Sun, M. et al. NuMA regulates mitotic spindle assembly, structural dynamics and function via phase separation. Nat Commun 12, 7157 (2021).

63. Huang, J., Liang, Z., Guan, C., Hua, S. & Jiang, K. WDR62 regulates spindle dynamics as an adaptor protein between TPX2/Aurora A and katanin. Journal of Cell Biology 220, e202007167 (2021).

64. Burns, K. M., Wagenbach, M., Wordeman, L. & Schriemer, D. C. Nucleotide Exchange in Dimeric MCAK Induces Longitudinal and Lateral Stress at Microtubule Ends to Support Depolymerization. Structure 22, 1173–1183 (2014).

65. Štimac, V., Koprivec, I., Manenica, M., Simunić, J. & Tolić, I. M. Augmin prevents merotelic attachments by promoting proper arrangement of bridging and kinetochore fibers. eLife 11, e83287 (2022).

66. Waters, J. C., Mitchison, T. J., Rieder, C. L. & Salmon, E. D. The kinetochore microtubule minus-end disassembly associated with poleward flux produces a force that can do work. MBoC 7, 1547–1558 (1996).

67. Desai, A., Maddox, P. S., Mitchison, T. J. & Salmon, E. D. Anaphase A Chromosome Movement and Poleward Spindle Microtubule Flux Occur At Similar Rates in Xenopus Extract Spindles. J Cell Biol 141, 703–713 (1998).

68. Sharp, D. J. & Rogers, G. C. A Kin I-Independent Pacman: Flux Mechanism for Anaphase A. Cell Cycle 3, 705–708 (2004).

69. Zhai, Y., Kronebusch, P. J. & Borisy, G. G. Kinetochore microtubule dynamics and the metaphase-anaphase transition. The Journal of cell biology 131, 721–734 (1995).

70. Brust-Mascher, I., Civelekoglu-Scholey, G., Kwon, M., Mogilner, A. & Scholey, J. M. Model for anaphase B: Role of three mitotic motors in a switch from poleward flux to spindle elongation. Proceedings of the National Academy of Sciences 101, 15938–15943 (2004).

71. Rath, U. et al. The Drosophila Kinesin-13, KLP59D, Impacts Pacman- and Flux-based Chromosome Movement. MBoC 20, 4696–4705 (2009).

72. Schelski, M. & Bradke, F. Microtubule retrograde flow retains neuronal polarization in a fluctuating state. Science Advances 8, eabo2336 (2022).

73. Burute, M., Jansen, K. I., Mihajlovic, M., Vermonden, T. & Kapitein, L. C. Local changes in microtubule network mobility instruct neuronal polarization and axon specification. Science Advances 8, eabo2343 (2022).

74. Ganem, N. J., Godinho, S. A. & Pellman, D. A mechanism linking extra centrosomes to chromosomal instability. Nature 460, 278–282 (2009).

75. Silkworth, W. T., Nardi, I. K., Scholl, L. M. & Cimini, D. Multipolar Spindle Pole Coalescence Is a Major Source of Kinetochore Mis-Attachment and Chromosome Mis-Segregation in Cancer Cells. PLOS ONE 4, e6564 (2009).

76. Gudlin, L. et al. A Universal Scaling Law for Mitotic Spindles Driven by Chromosome Crowding. 2025.03.05.641650 Preprint at 10.1101/2025.03.05.641650 (2025).

77. Harasymiw, L. A., Tank, D., McClellan, M., Panigrahy, N. & Gardner, M. K. Centromere mechanical maturation during mammalian cell mitosis. Nat Commun 10, 1761 (2019).

78. Dogterom, M. & Yurke, B. Measurement of the Force-Velocity Relation for Growing Microtubules. Science 278, 856–860 (1997).

79. Cameron, L. A. et al. Kinesin 5–independent poleward flux of kinetochore microtubules in PtK1 cells. J Cell Biol 173, 173–179 (2006).

80. Miyamoto, D. T., Perlman, Z. E., Burbank, K. S., Groen, A. C. & Mitchison, T. J. The kinesin Eg5 drives poleward microtubule flux in Xenopus laevis egg extract spindles. J Cell Biol 167, 813–818 (2004).

81. Garcia, M. A., Koonrugsa, N. & Toda, T. Two Kinesin-like Kin I Family Proteins in Fission Yeast Regulate the Establishment of Metaphase and the Onset of Anaphase A. Current Biology 12, 610–621 (2002).

82. West, R. R., Malmstrom, T. & McIntosh, J. R. Kinesins klp5(+) and klp6(+) are required for normal chromosome movement in mitosis. J Cell Sci 115, 931–940 (2002).

83. Gergely, Z. R., Crapo, A., Hough, L. E., McIntosh, J. R. & Betterton, M. D. Kinesin-8 effects on mitotic microtubule dynamics contribute to spindle function in fission yeast. MBoC 27, 3490–3514 (2016).

84. Toso, A. et al. Kinetochore-generated pushing forces separate centrosomes during bipolar spindle assembly. J Cell Biol 184, 365–372 (2009).

85. Tan, C. H. et al. The equatorial position of the metaphase plate ensures symmetric cell divisions. eLife 4, e05124 (2015).

86. Guerreiro, A. & Meraldi, P. AA344 and AA345 antibodies recognize the microtubule network in human cells by immunofluorescence. Antibody Reports e17–e17 (2019) doi:10.24450/journals/abrep.2019.e17.

